# *Staphylococcus epidermidis* Genomic Plasticity Modulates Horizontal Gene Transfer

**DOI:** 10.64898/2026.05.01.722213

**Authors:** My Tran, Angel Hernandez Viera, Jessica Rosu, Patricia Tran, Doran Goldman, Charlie Mo

## Abstract

Horizontal gene transfer (HGT) is an important driver of evolution, enabling organisms to rapidly adapt to environmental stressors. In bacteria, anti-mobile genetic element (MGE) defense systems are believed to limit HGT, raising the question of how bacterial populations balance defense with the acquisition of genetic novelty. Most insights into anti-MGE defense come from heterologous experimental setups, where candidate systems are artificially expressed in a host and observed over only a few generations. Such approaches can obscure the long-term evolutionary and ecological dynamics of native anti-MGE systems. Here, we show that laboratory-evolved *Staphylococcus epidermidis* populations rapidly delete a large, flexible genomic region containing anti-MGE defense systems and other accessory genes. This process generates stable, heterogeneous populations composed of cells that either retain or lack these defenses. Loss of this region is associated with a marked increase in HGT. Notably, the balance between anti-MGE defense and HGT is strongly influenced by environmental conditions. In nutrient-rich liquid media, defense systems and adjacent accessory genes are frequently lost, whereas under stress, such as antibiotic exposure, these defenses are maintained, leading to reduced HGT rates. This genetic and phenotypic heterogeneity enhances adaptive potential by enabling the uptake of beneficial MGEs while preserving protection under adverse conditions. Overall, the rapid gain and loss of anti-MGE defense systems highlight how environmental factors, independent of MGEs themselves, can shape HGT dynamics.

## Introduction

Horizontal gene transfer (HGT) is a fundamental biological process that enables the exchange of genetic material between organisms. In microbial communities, HGT plays a central role in evolution by shaping pangenomes and accelerating evolution; a canonical example is the dissemination of antibiotic resistance genes among bacterial pathogens. HGT is mediated by mobile genetic elements (MGEs), including bacteriophages (phages), plasmids, and transposons, which transfer genetic payloads that can enhance host fitness and survival. For instance, the lysogenic phage φCTx encodes toxins in *Vibrio cholerae* that increase bacterial virulence during infection (Waldor & Mekalanos, 1996). However, MGEs can also impose substantial costs on their hosts: parasitic plasmids and temperate phages may create metabolic burdens (Baltrus, 2013; Harrison & Brockhurst, 2017; San Millan & MacLean, 2017), while lytic phages can decimate bacterial populations through cell lysis (Salmond & Fineran, 2015). Thus, MGEs – and HGT more broadly – can be both beneficial and deleterious to bacteria (Haudiquet et al., 2022).

In response to the threat posed by parasitic MGEs, bacteria and archaea have evolved a wide arsenal of anti-MGE defense systems that detect and neutralize invading genetic material (Bernheim & Sorek, 2020). These systems are mechanistically diverse yet act as barriers to HGT between microbial populations (Bernheim & Sorek, 2020; Doron et al., 2018; Koonin et al., 2017; Rocha & Bikard, 2022). *In vitro* studies demonstrate that defense systems can inhibit the replication and propagation of virtually all known classes of MGEs, including lytic and temperate phages, plasmids, pathogenicity islands, and transposons (Beamud et al., 2024). Bioinformatic and metagenomic analyses support these conclusions at broader longitudinal scales. For example, the abundance of restriction–modification (RM) systems is inversely correlated with MGE frequency across bacterial taxa (J. Y. H. Lee et al., 2019a), while in marine *Vibrio* species, the presence of phage defense elements is anti-correlated with phage susceptibility (Hussain et al., 2021). Evolutionarily, these observations suggest that MGEs impose selective pressures that favor the maintenance of bacterial defense systems within bacterial populations (Hussain et al., 2021; Westra & Levin, 2020). However, most of our understanding of anti-MGE defense stems from heterologous experimental systems, where defense genes are expressed ectopically in a host bacterium and studied for only a few generations (Doron et al., 2018; Gao et al., 2020; Vassallo et al., 2022). As a result, how these defenses shape HGT across different ecologies and selective regimes is poorly understood and is a vibrant area of research (Gophna et al., 2015; Hussain et al., 2021; LeGault et al., 2021; Rocha & Bikard, 2022; Westra & Levin, 2020).

To investigate the dynamics of HGT and anti-MGE defense within a bacterial population, we performed experimental evolution of *Staphylococcus epidermidis*. A key advantage of laboratory evolution is the ability to analyze populations under precise experimental conditions, enabling one to determine causal links between environmental factors and evolutionary outcomes (Barrick & Lenski, 2013; Stroud & Ratcliff, 2025). *S. epidermidis* is both a ubiquitous commensal of the human skin and an opportunistic pathogen. In its commensal state, *S. epidermidis* protects the skin microbiome from pathogenic invasion and contributes to immune priming (Otto, 2009; Severn & Horswill, 2023). In contrast, during its pathogenic lifestyle, *S. epidermidis* is a leading cause of nosocomial and implant-associated infections (Otto, 2009). *S. epidermidis* possesses an open pangenome, with approximately 20% of its genome comprising variable genes (Brockhurst et al., 2019; Conlan et al., 2012; Doron et al., 2018; Takeuchi et al., 2005), indicative of extensive HGT both within *S. epidermidis* populations and between *S. epidermidis* and other species (Zhou et al., 2020). *S. epidermidis* genomes are also highly enriched in insertion sequence (IS) elements, which can mediate phase variation and rapid spontaneous genomic deletions and rearrangements (Schoenfelder et al., 2010). Thus, extensive HGT and genomic plasticity are thought to facilitate *S. epidermidis* adaptation to diverse ecological niches (Severn & Horswill, 2023). However, *S. epidermidis* strains also encode numerous anti-MGE defense systems targeting a broad range of MGEs. Metagenomic analyses indicate that individual strains can carry three to four distinct defense systems that tend to cluster downstream of *SCCmec* cassettes (Hossain et al., 2024). Indeed, these defense systems are thought to be the reason why many *S. epidermidis* strains are notoriously refractory to phage infection, plasmid uptake and genetic manipulation (Costa et al., 2017; J. Y. H. Lee et al., 2019a). Together, these observations raise a central question: how do the levels of anti-MGE defense and HGT change in evolving *S. epidermidis* populations?

Here, we report that *S. epidermidis* with anti-MGE defenses can rapidly evolve higher rates of HGT and modulate gene flow in response to selective pressure. We performed laboratory evolution on *S. epidermidis* RP62a (RP62a), which is a strain that carries four distinct defense systems and is highly resistant to conjugation, transformation, and phage infection. After only 80 generations, conjugation, transformation, and phage plaquing all increased by at least three orders of magnitude. This increase in HGT is accompanied by a reduction in the RP62a genome by approximately 320 kB, which is greater than 10 percent of the total genome (∼2.6 Mbp). The genomic loss occurs in a flexible genomic region that is flanked by short insertion (IS) elements and replete with accessory genes, including the four defense systems and genes mediating drug resistance and virulence. However, a subpopulation of RP62a with the complete accessory genes persisted, forming stable heterogeneous populations. These evolved *S. epidermidis* populations have increased acquisition of MGEs, thereby enhancing the population’s adaptability, with the trade-off cost of increased susceptibility to lytic phage infections. Subjecting bacterial populations to selective pressures, such as antibiotic treatment, stabilized the flexible region, causing anti-MGE defense to be maintained, and reducing HGT levels. This linkage of anti-MGE defense with other accessory functions, such as drug resistance and biofilm production, places defense systems and HGT under the control of diverse selective pressures and environments.

## Results

### Permissiveness of HGT rapidly increases in evolving *Staphylococcus epidermidis* populations

*S. epidermidis* RP62a (RP62a) is a pathogenic strain originally isolated from a catheter-associated infection in the 1970s (Supplemental Table 1) (Gill et al., 2005). RP62a is capable of robust biofilm formation and harbors a type II *SCCmec* element, which confers resistance to β-lactam antibiotics (Datta et al., 2021). Downstream of *SCCmec*, RP62a encodes four anti-MGE defense systems: a type III CRISPR-Cas system; a eukaryotic-like serine/threonine kinase (Stk2); a nuclease–helicase (Nhi); and a type I restriction–modification (RM) system (Bari et al., 2022; Depardieu et al., 2016; J. Y. H. Lee et al., 2019a; Marraffini & Sontheimer, 2010). Genomically, these four systems are clustered within approximately 30 kB of one another, forming a so-called defense island (Bernheim & Sorek, 2020) (Figure 1A). Collectively, these defenses restrict infection by diverse MGEs. Stk2 and Nhi inhibit bacteriophage infection, while the first spacer of the CRISPR-Cas system targets the highly conserved *nickase* (*nes*) gene of the staphylococcal plasmid pG0400, which belongs to the pSK41/pGO1 family of multi-resistance plasmids capable of promoting the conjugative transfer of resistance genes (Bari et al., 2022; Depardieu et al., 2016; Marraffini & Sontheimer, 2008). The type I RM system also provides additional protection against plasmids and phages (Costa et al., 2017). Previous studies have estimated that RP62a populations can spontaneously lose defense systems (Hossain et al., 2024; Jiang et al., 2013), yet how HGT rates change over multiple generations – and how environmental conditions modulate these dynamics – remains unknown. Using RP62a as an experimental model, we asked how HGT rates evolve in defense system–positive bacterial populations over extended evolutionary timescales. Importantly, our experimental evolution system enabled us to systematically test the effects of different selective pressures on *S. epidermidis* HGT.

**Figure 1:**
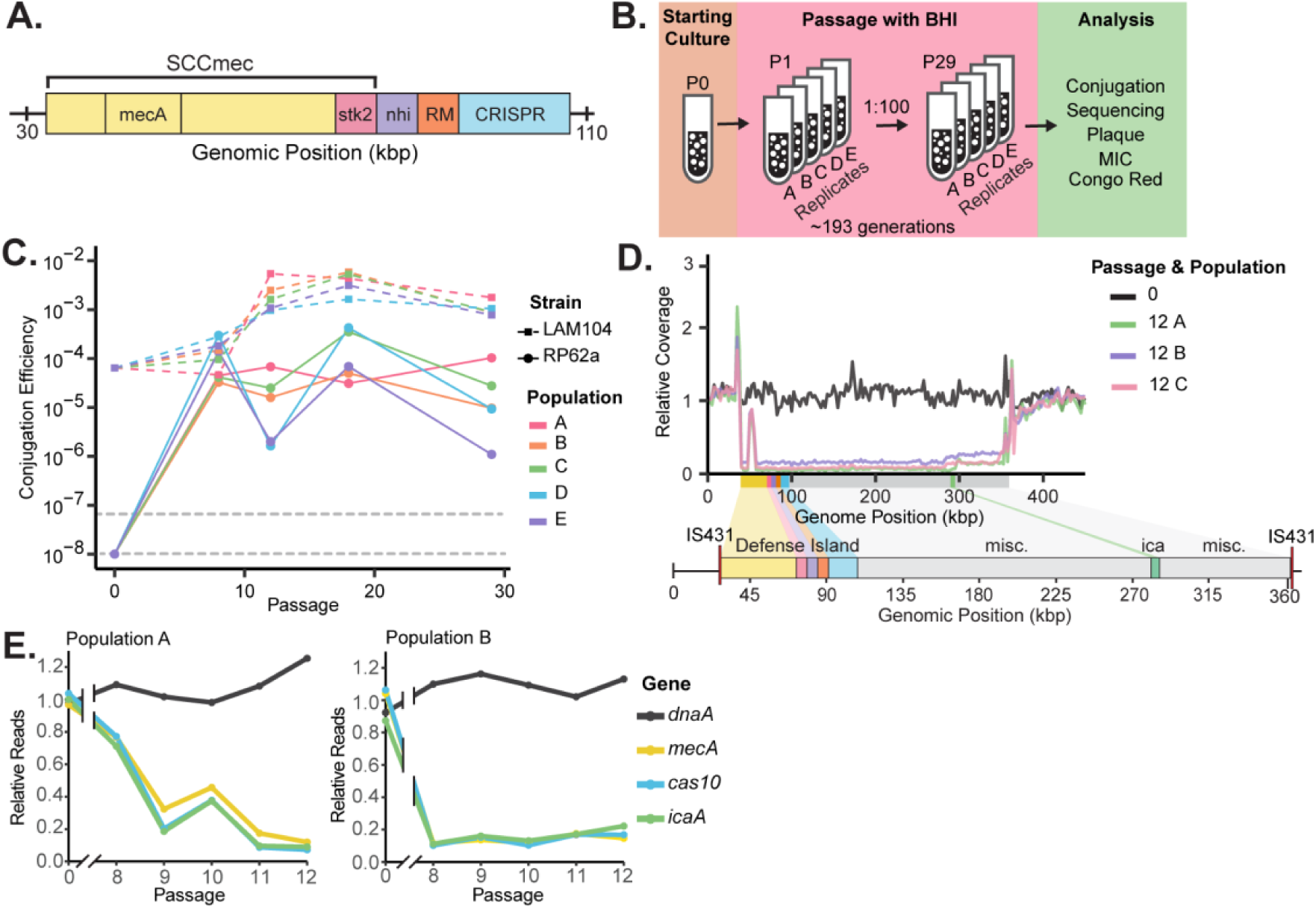
Evolved RP62a Populations have increased HGT due to genomic deletion. **A.** Schematic of the anti-MGE defense island and *SCCmec* in *S. epidermidis* RP62a. **B.** Experimental evolution of *S. epidermidis* RP62a. Starting from a parental clone (P0), 5 independent cultures (A-E) were passaged every 24 hours for 29 days (P29). During passaging, the cultures were sequenced and subjected to phenotypic analyses. **C.** Conjugation efficiency of evolving *S. epidermidis* RP62a and LAM104 populations. *S. epidermidis* populations at passages 0, 8, 12, 18, and 29 were conjugated with the staphylococcal conjugative plasmid, pG0400. Conjugation efficiency was measured by dividing transconjugants by the total number of *S. epidermidis* recipients. The two dotted lines represent the upper and lower bound limits of detection. **D**. Short-read coverage plot of *S. epidermidis* RP62a P0 and three independent P12 populations (A-C). Shown are the first 450 kbp of the 2.6 Mbp *S. epidermidis* RP62a genome. The y-axis represents the reads per million (RPM) normalized to the RPM of *dnaA,* a housekeeping gene. The colored region beneath the coverage plot is a schematic of the genes present in the region, which include *SCCmec*, the anti-MGE defense island, the *ica* locus, and other miscellaneous genes The red line flanking the accessory genes represents copies of the insertion element IS431. **E.** Relative reads measuring the presence of a gene in the population through passages. Relative reads were measured by dividing the average number of reads at a gene locus by the average number of total reads. The relative reads were calculated across P0, P8-P12. Replicates A and B are shown.

We passaged five independent cultures of RP62a for 29 days (Figure 1B) in rich Brain Heart Infusion (BHI) medium. The cultures were founded from an individual RP62a clone, and at each daily passage, 1% of the previous culture was transferred into fresh medium. At defined time points, we assayed the ability of these evolved populations to acquire the plasmid pG0400, which carries mupirocin resistance. In parallel, we passaged five populations of LAM104, a spacer-negative (ΔCRISPR) derivative that cannot target pG0400 (Supplemental Table 1) (Marraffini & Sontheimer 2008). Consistent with prior reports, conjugation was not detectable in the parental (P0) RP62a strain above the limit of detection (∼10⁻⁸ transconjugants per recipient cell), indicative of active CRISPR–Cas immunity (Figure 1C) (Marraffini & Sontheimer, 2008). In contrast, LAM104 P0 populations exhibited conjugation frequencies ranging from 10^-5^ and 10^-4^.

Over the course of passaging, we found that the conjugation efficiency increased rapidly in all five RP62a populations, reaching frequencies >10⁻⁵ by passage 8 (P8) in every population (Figure 1C). In some lineages (Populations D and E), conjugation frequencies fluctuated at later passages but consistently remained above 10⁻⁶. We also observed an increase in the five LAM104 populations: by P12, pG0400 conjugation frequencies increased from ∼10⁻⁴ to ∼10⁻³, indicating an approximately tenfold enhancement even in the absence of CRISPR targeting. Together, these evolution experiments demonstrate that pG0400 conjugation frequencies in *S. epidermidis* populations can increase substantially over a relatively low number of bacterial generations (∼80 generations).

### Evolved RP62a populations display a loss in frequency of a flexible genomic region

To identify the genetic basis underlying the increased plasmid conjugation rates, we performed whole-genome, short-read sequencing on three of the five passaged *S. epidermidis* RP62a populations (A–C) to determine genomic changes. By passage 12, all sequenced populations exhibited a loss of read coverage across a genomic region downstream of the *oriC,* flanked by two copies of the insertion sequence IS431 in the same orientation (Figure 1D; Supplemental Figure 2)(Wada et al., 1991). This deletion spans approximately 320 kb and removes numerous accessory genes, including all four anti-MGE defense systems, the *SCCme*c cassette, and the *ica* locus responsible for polysaccharide intercellular adhesin (PIA) synthesis (Table S2). Such large-scale deletions are not unique to the RP62a strains, as they have previously been reported in other disease-associated *S. epidermidis* isolates (Schoenfelder et al., 2010).

We further sequenced individual clones isolated from passage 12 of Population A and found that all clones harbored the ∼320 kb deletion (Supplemental Figure 1). Moreover, whole-genome sequencing of four P12 LAM104 populations (A–D) revealed the same ∼320 kb deletion in each lineage (Supplemental Figure 2A & B), paralleling the genomic changes observed in evolved RP62a populations. The 10-fold enhanced conjugation efficiency observed in these ΔCRISPR populations is likely attributable to the loss of the remaining three defense systems, particularly the RM system, which is known to target incoming plasmid DNA (Costa et al., 2017).

To analyze the deletion event at a higher temporal resolution, we sequenced passages 8-12 (P8–P12) of RP62a Populations A and B. We quantified the ratio of mean read depth at representative loci relative to total genomic coverage. We examined genes located within the deleted region – *mecA* (the signature penicillin binding protein in *SCCmec*), *cas10* (the signature gene of the type III CRISPR-Cas system), and *icaA* (within the *ica* locus responsible for exopolysaccharide synthesis) – as well as *dnaA*, a housekeeping gene that initiates DNA replication, as an internal control (Gill et al., 2005; Marraffini & Sontheimer, 2008; McKenney et al., 1998; Menikpurage et al., 2021) Across passages, the mean coverage for *dnaA* remained stable at ∼100%, whereas read depth for *mecA*, *cas10*, and *icaA* declined, reaching ∼10% by P12 (Figure 1E). Population A exhibited a gradual decrease in coverage from P8 to P12, whereas Population B already showed markedly reduced coverage by P8, indicating that the deletion arose earlier in this lineage. These distinct trajectories suggest stochasticity in the timing of IS431-mediated loss of the ∼320 kb genomic region across independently evolving populations (Figure 1E).

We further extended this analysis to later passages (P18 and P29) in Population A. The relative read depth of the accessory genes remained below 20% at P18 and P29, suggesting that deletion-bearing cells did not fully sweep the population: a consistent minority retained the intact ∼320 kb region at P12, P18, and P29 (Figure 1E; Supplemental Figure 2C). These results indicate that clonal RP62a populations rapidly evolve into heterogeneous communities comprising at least two coexisting genotypes – one lacking the accessory gene region (Δaccessory) and one retaining the wild-type genome. Consistent with previous work, competition assays suggest that carriage of the flexible genomic region does not impose a strong fitness cost or benefit on *S. epidermidis* RP62a (Supplemental Table 1, Supplemental Figure 3A) (Jiang et al., 2013). These genotypes persist at relatively stable proportions over multiple passages, suggesting a balance between the fitness advantages and costs associated with loss of the accessory region. Notably, our results in *S. epidermidis* are consistent with reports of similar diversification events observed in the *oriC* environ of other staphylococci (Bouchami et al., 2023; Takeuchi et al., 2005). We therefore speculate that our findings in RP62a could be broadly applicable across different staphylococci (Takeuchi et al., 2005).

Motivated by our observations that the variable region is flanked by two copies of the insertion sequence IS431, we asked whether anti-MGE defense systems in RP62a – and more broadly across *S. epidermidis* – are commonly associated with insertion sequences in general. Across different bacterial species, defense systems are frequently associated with MGE, enabling the gain or loss of anti-MGE phenotypes independent of the host core genome (Delaney et al., 2012). We retrieved 201 complete *S. epidermidis* genomes from NCBI and used DefenseFinder to identify anti-MGE defense genes (Néron et al., 2023; Tesson et al., 2022). In agreement with prior reports, anti-MGE systems were enriched in a genomic region downstream of *SCCmec* and the *rlmH* gene, which encodes for ribosomal RNA large subunit methyltransferase H (Hossain et al., 2024). In parallel, we identified insertion sequences using ISfinder and ISEScan (Siguier, 2006; Xie & Tang, 2017). Strikingly, anti-MGE systems consistently colocalized with multiple insertion sequences within the first ∼150 kb of *S. epidermidis* genomes. Together, these analyses suggest that the high local density of short transposable elements near anti-MGE defense loci can create recombination hotspots that facilitate recurrent deletion of large accessory regions across *S. epidermidis* lineages (Supplemental Figure 4).

### Evolved RP62A populations show increased permissiveness to different classes of MGEs

While the CRISPR–Cas system in RP62a provides sequence-specific protection against plasmids carrying the *nes* target, the broader defense island encodes multiple systems that together can restrict diverse classes of MGEs. We therefore hypothesized that loss of the defense island through genomic deletion would increase permissiveness to multiple MGEs.

To test this prediction, we first performed additional conjugation assays using pG0400mut, a derivative of pG0400 containing silent mutations in the *nes* target that render it invisible to CRISPR targeting (Figure 2A) (Jiang et al., 2013). In P0 RP62a, pG0400mut conjugated at a frequency of ∼1 × 10⁻⁴, approximately three orders of magnitude higher than wild-type pG0400. In the evolved P12 populations, the conjugation frequency of pG0400mut significantly increased by an additional order of magnitude across all five lineages, reaching ∼1×10⁻³ (Figure 2A). These rates matched those observed in LAM104 and LM1680. These results further indicate that defense systems outside of CRISPR-Cas contribute substantially to the restriction of plasmid conjugation in RP62a (J. Y. H. Lee et al., 2019b; Mercer & Loutit, 1979).

**Figure 2:**
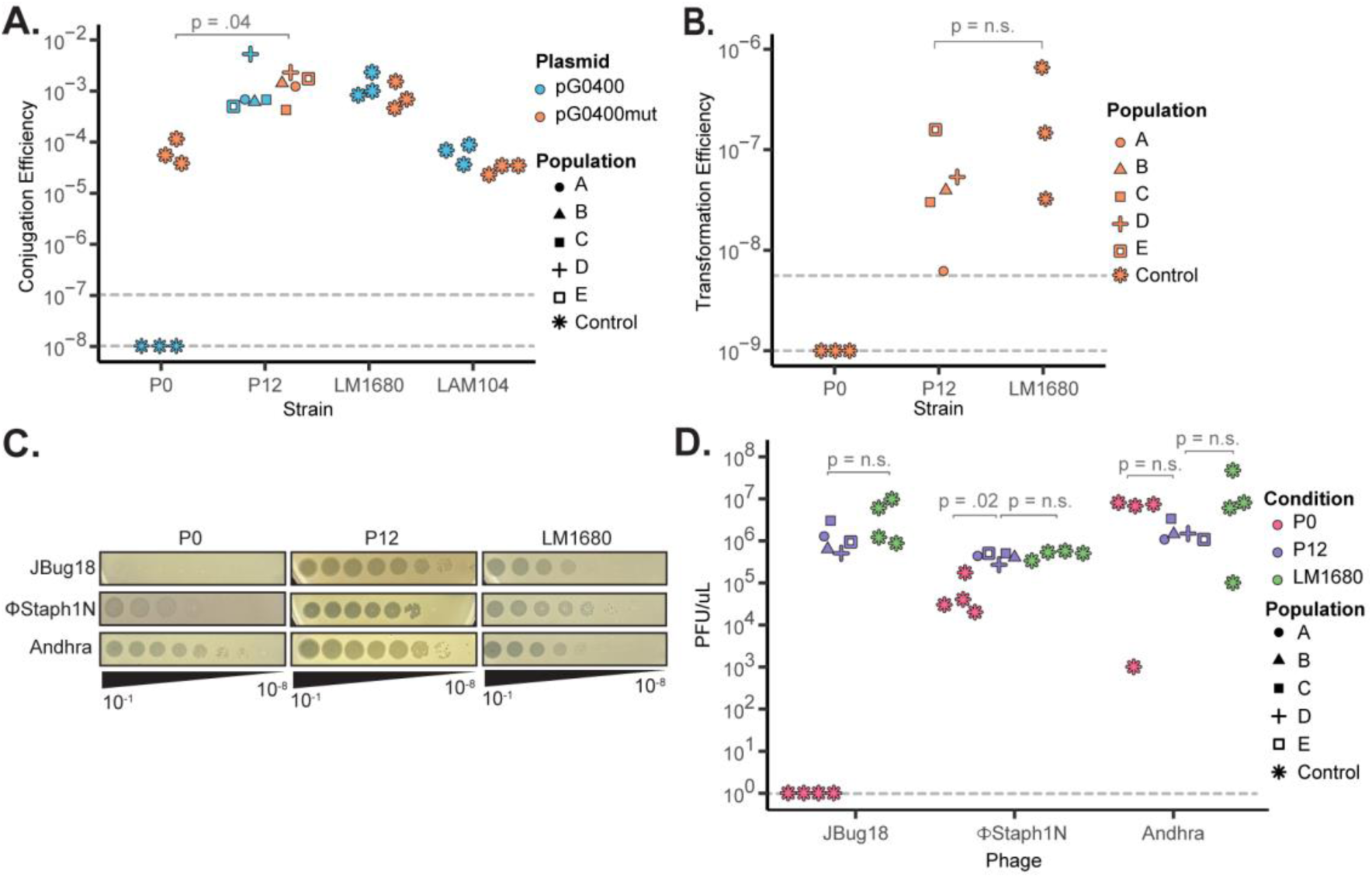
Evolved *S. epidermidis* populations display increased permissiveness to HGT. Experiments were performed on parental (P0) and evolved (P12) RP62a populations. LM1680 denotes a clonal Δaccessory strain lacking anti-MGE defense, while LAM104 is a strain lacking CRISPR immunity. **A.** Conjugation efficiencies of *S. epidermidis* populations with plasmids pG0400 or pG0400mut. **B.** Transformation efficiencies of *S. epidermidis* populations with pC194. **C.** Viral plaquing on *S. epidermidis* populations. Each population was challenged with a 10-fold dilution series of phages JBug18, Andhra, and ΦStaph1N (1 x 10^-1^ – 1 x 10^-8^). The phage titer was calculated as PFU/µL. **D.** Plaquing efficiencies of JBug18, Andhra, and ΦStaph1N on populations of *S. epidermidis*. For Panels A, C, and D: shapes indicate replicate P12 populations (A-E); where applicable, two dashed lines represent the upper and lower bound limits of detection; statistical significance was calculated using a two-sided Mann-Whitney Test.

We next assessed whether loss of the defense island could affect plasmid transformation. *S epidermidis* RM systems are a strong barrier against plasmid transformation, making isolates highly recalcitrant to electroporation (Costa et al., 2017). To assess if the loss of the RM system increases transformation efficiency, we generated electrocompetent cells from parental P0 and evolved P12 populations. We measured the transformation efficiency of pC194, a staphylococcal plasmid conferring chloramphenicol resistance (Horinouchi & Weisblum, 1982). In P0 RP62a, transformants were not detected above the limit of detection (∼10⁻⁹; Figure 2B). In contrast, RP62a P12 populations exhibited a marked increase in transformation efficiency, ranging from ∼10⁻⁸ to 10⁻⁷. These levels are comparable to those observed in LM1680. These findings demonstrate that the evolutionary loss of the defense island renders RP62a more broadly permissive to plasmid acquisition, extending beyond CRISPR-targeted conjugation to transformation.

Lastly, we assessed whether the evolutionary loss of the defense island also altered resistance to lytic phage infection. We challenged RP62a P12 populations with three staphylococcal phages: JBug18, ΦStaph1N, and Andhra (Figure 2C). All three phages efficiently infect the LM1680 strain but exhibit distinct infectivity profiles against parental P0 RP62a: the Nhi defense system, which is part of the defense island, confers resistance against JBug18, resulting in highly reduced plaquing, while the defense systems does not target Andhra or ΦStaph1N, permitting both phages to plaque on RP62a (Bari et al., 2022). Relative to P0 RP62a, all P12 populations displayed markedly increased phage susceptibility, with plaquing efficiencies increasing by more than six orders of magnitude for JBug18, and by one order of magnitude for ΦStaph1N in P12 populations compared to P0 (Figure 2D). Susceptibility to Andhra remained high across both P0 and P12 (Figure 2D).

### Evolved RP62A cells show altered virulence and drug resistance phenotypes

In addition to the defense island, the flexible genomic region encodes several genes implicated in *S. epidermidis* pathogenesis. The *SCCmec* element confers resistance to β-lactams, while *ica* produces PIA, which promotes biofilm formation and immune evasion (Paharik & Horswill, 2016). We therefore assessed whether the evolved populations retained resistance to the β-lactam oxacillin and the capacity to produce PIA.

RP62a P0 populations exhibited high-level oxacillin resistance, with minimal inhibitory concentrations (MICs) of ∼64–128 µg/mL, whereas LM1680 was highly sensitive, with an MIC of ∼0.75 µg/mL (Fig. 3A). Meanwhile, RP62a P12 populations displayed a heterogeneous resistance phenotype, with a portion of cells showing higher level of oxacillin sensitivity than others (Fig. 3A). To further quantify this observation, we measured MICs using broth dilution. Consistent with the visual assay, RP62a P0 and LM1680 exhibited MICs of >64 µg/mL and <1 µg/mL, respectively, while RP62a P12 populations showed a modest reduction in resistance, with MICs of ∼32–64 µg/mL (Figure 3B). These data indicate that, despite heterogeneity, evolved P12 RP62a populations retain substantial β-lactam resistance.

**Figure 3.**
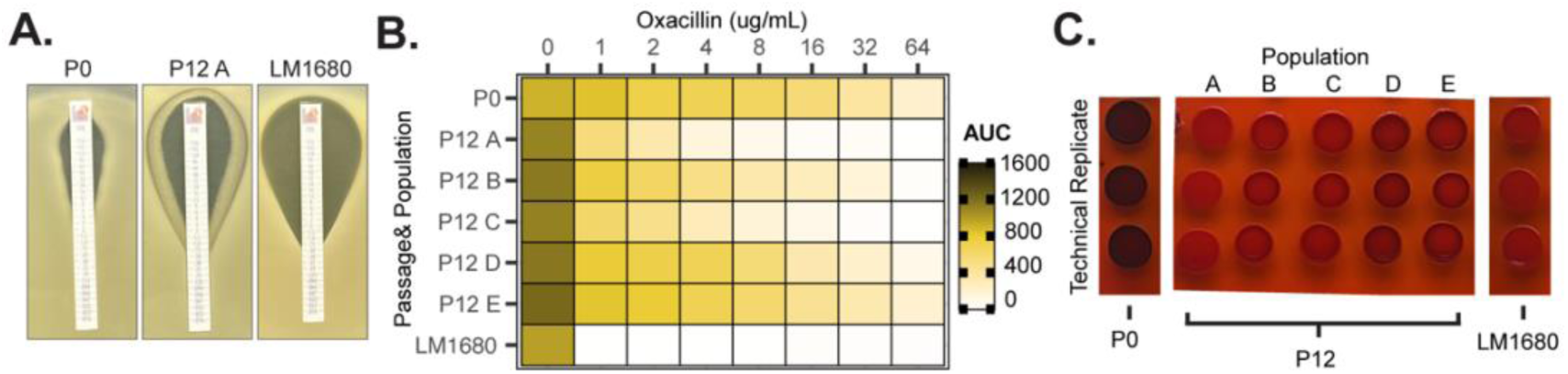
β-lactam resistance and <u>P</u>olysaccharide Intercellular <u>A</u>dhesin (PIA) production in evolved *S. epidermidis* populations. Experiments were performed on parental (P0) and evolved (P12) RP62a populations. LM1680 denotes a clonal Δaccessory strain lacking *ica.* **A.** Representative image of oxacillin minimum inhibitory concentration (MIC) of *S. epidermidis* populations. Shown is a plate from population P12 A. **B.** Oxacillin MICs shown as an area under the curve (AUC) heatmap of the *S. epidermidis* populations. A-E indicate independent biological populations of the P12 populations. **C.** Congo red assay on *S. epidermidis* populations. A-E indicate independent biological populations, each plated in three technical populations. Black coloration is PIA+, red coloration is PIA-.

We next examined PIA production using Congo Red agar, which distinguishes PIA-producing [PIA(+)] colonies (black) from non-producing [PIA(−)] colonies (red) (Kaiser et al., 2013; J.-S. Lee et al., 2016). RP62a P0 appeared uniformly black, whereas RP62a P12 populations displayed red coloration, more similar to LM1680 (Fig. 3C). This shift indicates a marked reduction in PIA production following evolution.

Beyond *SCCmec* and *ica*, the deleted region also harbors numerous other accessory genes. Although some have predicted functions (e.g., arsenic resistance), most remain uncharacterized (Supplemental Table 2). Altogether, our findings demonstrate that RP62a can lose more than 10% of its genome, including all known anti-MGE defense systems, within fewer than 80 generations. This large-scale genomic deletion drives an increase in HGT while simultaneously modulating drug resistance and virulence phenotypes of the populations.

### Antibiotic selection and sessile growth modulate heterogeneity and HGT levels in *S. epidermidis* populations

We next asked whether environmental stressors could influence the retention of anti-MGE defense systems in evolving RP62a populations, thus modulating HGT rates. Because the defense island and adjacent accessory genes are consistently lost as a single unit, we hypothesized that selection acting on one locus could indirectly alter the frequency of linked genes within this region. In particular, we focused on *SCCmec* and *ica,* which are separated by ∼200 kb, are subject to distinct selective pressures, and encode key virulence determinants of RP62a (Severn & Horswill, 2023).

To test this idea, we passaged RP62a for 15 passages under two different selective conditions (Figure 4A). First, we passaged RP62a in liquid media at increasing oxacillin concentrations (3 µg/mL, 6 µg/mL, 12 µg/mL), directly selecting for the *SCCmec*-encoded β-lactam resistance. To enrich for intact *ica*, we passaged RP62a under sessile (SP) conditions using polystyrene culture flasks, a regimen that favors the formation of biofilm with adhering bacteria (Ziebuhr et al., 1999). We then compared the genotypes and phenotypes of these populations with those of the control RP62a populations passaged in parallel in BHI medium (Figure 4B, Supplemental Table 3).

**Figure 4.**
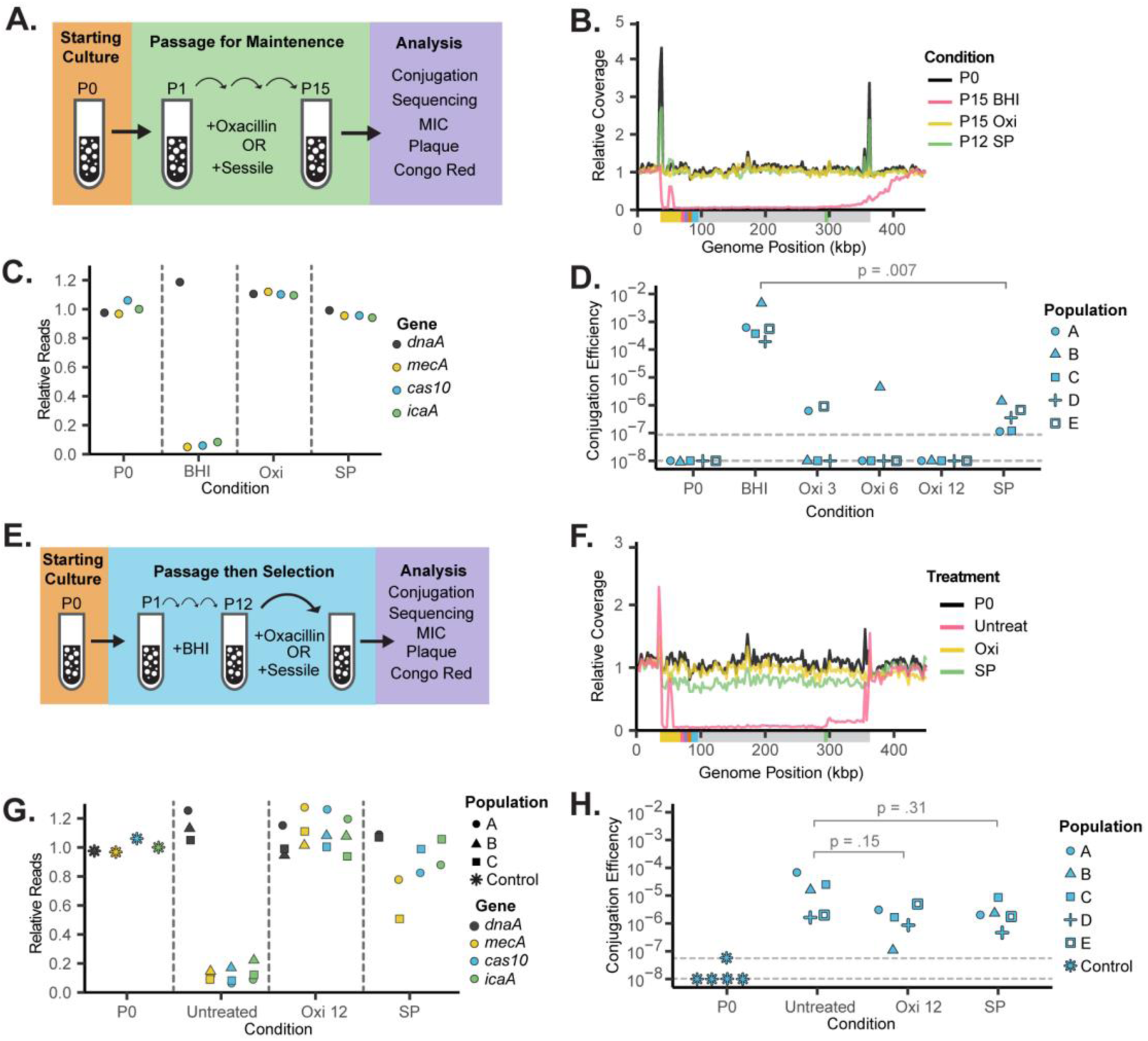
Antibiotic selection and sessile growth modulate anti-MGE defense and HGT in *S. epidermidis*. **A.** RP62a populations were passaged in the presence of oxacillin or under sessile conditions (SP). **B.** Short-read coverage plot of passaged *S. epidermidis* populations. Shown are the first 450 kbps out of 2.6 Mbp of the RP62a genome. The y-axis shows the reads-per-million (RPM), normalized to the RPM at the *dnaA* position. The colored region beneath the coverage plot is a schematic of the genes present in the region, as seen in Figure 1D. BHI: non-selective; SP: sessile conditions; Oxi: 3 µg/mL oxacillin. **C.** Relative abundance of *dnaA, mecA, cas10,* and *icaA* from the sequencing data shown in Panel B. **D.** pG0400 conjugation efficiency of *S. epidermidis* populations grown under different conditions. The concentration of oxacillin (µg/mL) is indicated next to the oxacillin abbreviation. The 5 evolving populations are indicated by the different symbols. The two dotted lines represent the upper and lower bound limits of detection. **E.** Treatment of evolved RP62a with 12 ug/mL of oxacillin (Oxi 12) or sessile passaging (SP). **F.** Short-read coverage plot of treated and untreated *S. epidermidis* populations. Shown are the first 450 kbps out of 2.6 Mbp of the RP62a genome. The y-axis shows the reads-per-million (RPM), normalized to the RPM at the *dnaA* position. The colored region beneath the coverage plot is a schematic of the genes present in the region, as seen in Figure 1D. SP: sessile conditions; Oxi: 12 µg/mL oxacillin. **G.** Relative abundance of *dnaA, mecA, cas10,* and *icaA* from the sequencing data shown in Panel F**. H.** pG0400 conjugation efficiency of treated and untreated *S. epidermidis* populations. Independent Populations are indicated by shapes. Statistical significance was calculated using a two-sided Mann-Whitney Test. The two dotted lines represent the upper and lower bound limits of detection.

Unlike BHI-passaged populations, populations that underwent SP or passaging in 3 µg/mL oxacillin did not produce the characteristic drop in coverage across the deleted region (Fig. 4B). Read depths at *mecA*, *cas10*, and *icaA* matched those of *dnaA* (Fig. 4C), confirming preservation of the defense island and surrounding accessory genes. Oxacillin-passaged and SP populations also displayed phenotypes comparable to those of unevolved P0 populations. In the absence of oxacillin, P15 populations showed elevated conjugation frequencies (∼10⁻²–10⁻³), consistent with the evolution experiment shown in Figure 1 (Fig. 4D). By contrast, oxacillin-evolved populations showed reduced levels of conjugation. At 12 µg/mL oxacillin, none of the five replicates exhibited detectable transfer above the assay limit (∼10⁻⁸), comparable to P0. Populations evolved in 6 or 3 µg/mL oxacillin showed limited conjugation, with only one and two populations, respectively, displaying detectable transfer at frequencies below 10⁻⁵. SP-evolved populations displayed markedly reduced conjugation (∼10⁻⁷–10⁻⁶) as well, significantly lower than BHI-passaged controls (Fig. 4D). Furthermore, consistent with their intact accessory regions, oxacillin-evolved populations and SP populations remained JBug18-resistant, PIA-positive, and displayed comparable oxacillin resistance to P0 populations (Supplemental Fig. 5).

Together, these phenotypic and genomic data show that selection on either *SCCmec* or *ica* is sufficient to stabilize the entire linked accessory region, including anti-MGE defense systems, and thereby suppressing HGT. Low, but detectable, conjugation in populations evolved under weaker oxacillin concentrations or SP. More broadly, these results show that non-MGE selective pressures can modulate the frequency of anti-MGE defense systems in evolving *S. epidermidis* populations.

We then asked whether reapplying a single selective pressure to evolved RP62a P12 populations could restore accessory gene frequencies at the population level (Figure 4E). Because P12 populations contain a minority of wild-type cells retaining the accessory region, we hypothesized that targeted selection would enrich for this subpopulation. We therefore treated P12 populations with 12 µg/mL oxacillin or subjected them to SP for nine additional passages, followed by genotypic and phenotypic characterization (Figure 4E, Supplemental Table 3).

In both oxacillin- and SP-treated populations, sequencing coverage increased across the ∼320-kb region (Fig. 4F; Supplemental Fig. 6A). Oxacillin-treated populations recovered *mecA*, *cas10*, and *icaA* to ∼80–100% frequency (Fig. 4G); SP-treated P12 populations showed near-complete recovery of *cas10* and *icaA*, while *mecA* frequencies varied between replicates (∼40% vs. ∼80%). Phenotypically, both treatments retained oxacillin resistance, increased the PIA(+) phenotype, and preserved JBug18 resistance (Supplemental Figure 6B-D). However, populations from both treatments exhibited only a modest, statistically insignificant, ∼10-fold decrease in conjugation relative to untreated P12 populations (Figure 4H). We isolated pG0400-positive clones and performed genotypic analysis of these transconjugant isolates. Sequencing of clones revealed the same large genomic deletion that includes the anti-MGE defense systems, *SCCmec*, and *ica* locus described previously. One isolate was an exception: a clone from the SP-treated populations (SP C2) retained the entire accessory region (Supplemental Figure 6E). However, the SP2 C2 clone had a frameshift mutation in the *csm2* gene of the type III CRISPR system, which would inactivate CRISPR targeting against pG0400. These findings suggest that treating an evolved population with oxacillin or exposing it to a biofilm-inducing environment will select for cells with the intact region but may not completely restore anti-MGE function. Remaining Δaccessory clones or mutants with defects in anti-MGE genes (e.g. SP C2) can still enable permissiveness to MGEs.

Lastly, we investigated how phage selection affects the evolved P12 populations. Notably, infection with JBug18 did not favor the recovery of the defense island or the surrounding accessory genes in the bacterial populations (Supplemental Figure 6A & G). JBug18-treated P12 displayed relatively high conjugation levels, oxacillin resistance, and variable PIA production (Supplemental Figure 6). However, JBug18-treated populations did become resistant to JBug18 and Andhra, but not ΦStaph1N (Supplemental Figure 6B). We also treated the P12 populations with either ΦStaph1N or Andhra. In both Andhra and Staph1N infections, surviving *S. epidermidis* populations exhibited a marked reduced ability to regrow in liquid culture. We were therefore unable to perform phenotypic and genotypic analysis on these populations. We note that these results do not preclude the possibility that predation by a different phage could help maintain the flexible genomic region and anti-MGE defense. Rather, as a whole, these selection results show that non-phage selective pressures, such as antibiotic treatment, can maintain antiphage defense within the bacterial population, thereby modulating HGT rates.

### Increased permissiveness to MGEs improves the adaptability of the bacterial population

HGT accelerates bacterial adaptation and evolution by rapidly distributing genes across populations (Woods et al., 2020). We hypothesized that the increased capacity for acquiring genetic novelty through HGT observed in evolved P12 populations would increase the populations’ adaptability relative to homogeneous wild-type or Δaccessory backgrounds. To test this, we conjugated pG0400 – which confers mupirocin resistance – into three clonal RP62a P0 populations, five heterogeneous RP62a P12 populations, and three clonal LM1680 populations. We then took these conjugated populations and passaged them for two rounds (R1→R2) in either non-selective BHI media (R1_mock_→R2_mock_) or in alternating treatments of two antibiotics, oxacillin and mupirocin (R1_oxi_→R2_mup_ or R1_mup_→R2_oxi_). For each passage, we quantified their ability to grow in the presence of oxacillin or mupirocin (Figure 5A).

**Figure 5:**
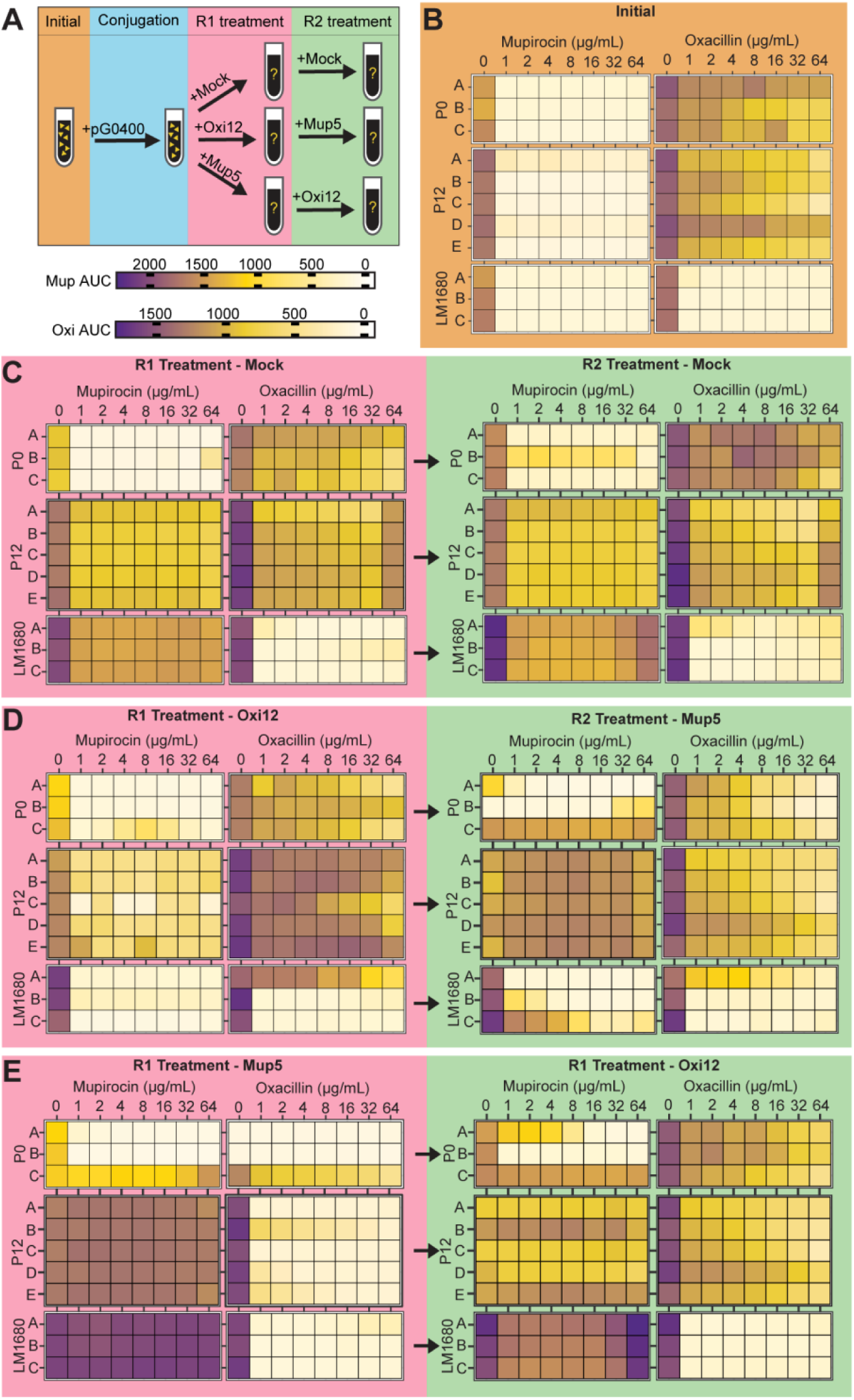
Increased susceptibility to HGT allows for higher adaptability in evolved populations. **A.** Three RP62a P0 populations, the 5 evolved P12 populations, and three LM1680 clones were conjugated with plasmid pG0400 and subjected to two rounds (R1, R2) of non-selective (mock) or alternating antibiotic treatment (Oxi12; Mup5). After each round, populations were tested for their ability to grow in increasing levels (µg/mL) of oxacillin and mupirocin, using area under the curve (AUC) measurements. **B.** Growth of *S. epidermidis* populations in oxacillin and mupirocin prior to pG0400 conjugation. **C-E.** Growth in oxacillin and mupirocin of the *S. epidermidis* populations following pG0400 conjugation and **C.** R1_mock_→R2_mock_; **D.** R1_oxi_→R2_mup_ treatment; and **E.** R1_mup_→R2_oxi_ treatment.

As expected, prior to pG0400 conjugation, RP62a P0 and P12 were mupirocin-sensitive and oxacillin-insensitive, whereas LM1680 was sensitive to both antibiotics (Figure 5B). Following pG0400 conjugation and non-selective passaging (R1_mock_→R2_mock_), RP62a P0 remained largely mupirocin-sensitive, while P12 and LM1680 showed increased ability to grow in mupirocin, consistent with higher pG0400 acquisition and retention when anti-MGE defenses are reduced or absent (Figure 5C). Oxacillin phenotypes were unchanged: RP62a P0 and P12 grow in the presence of oxacillin, while LM1680 showed highly reduced growth. Thus, the heterogeneous P12 populations readily achieved dual resistance (oxacillin via *SCCmec*; mupirocin via pG0400).

We next tested whether alternating antibiotic selection reshapes these composite sensitivity profiles (Figures 5D and E). Because anti-MGE defenses limit co-residence of *SCCmec* and pG0400 within single cells, we predicted that conjugated populations would comprise mixtures of oxacillin- and mupirocin-resistant subpopulations and that selection for one trait could purge the other from the population. Under oxacillin-then-mupirocin selection (R1_oxi_→R2_mup_), conjugated RP62a P0 populations remained predominantly oxacillin-insensitive and mupirocin-sensitive, with a single replicate gaining mupirocin resistance after R2_mup_ (Figure 5D). In LM1680, oxacillin selection likely depleted pG0400-bearing cells, preventing subsequent mupirocin enrichment; one replicate acquired oxacillin insensitivity. In contrast, all P12 populations retained resistance to both antibiotics, indicating that oxacillin selection did not eliminate the mupirocin-resistant fraction. We interpret this stability as a population-level effect: *SCCmec*(+) cells can buffer Δaccessory pG0400(+) cells, allowing mupirocin resistance to persist despite oxacillin selection.

Reversing the order (R1_mup_→R2_oxi_) yielded a complementary pattern (Figure 5E). RP62a P0 again remained largely mupirocin-sensitive (with Replicate C resistant), and transient oxacillin sensitivity after R1_mup_ was reversed by R2_oxi_. LM1680 remained mupirocin-resistant yet oxacillin-sensitive across rounds. P12 populations maintained mupirocin resistance throughout but exhibited a transient reduction in oxacillin resistance after R1_mup_, followed by recovery under R2_oxi_, consistent with selection-driven shifts in subpopulation frequencies rather than loss of either determinant.

Together, these results show that evolved P12 populations readily acquire MGEs and retain them under fluctuating selection, generating adaptive flexibility that homogeneous backgrounds lack. While antibiotic order modulates the relative abundance of resistant subpopulations, community-level buffering can stabilize partitioned traits, preserving dual resistance at the population level. By contrast, RP62a P0 is constrained by restrictive anti-MGE defenses, and LM1680 lacks key accessory functions, leaving both dependent on other routes for adaptation.

### Extended passaging results in diverse genomic variants

Bacterial populations are continuously evolving. We therefore asked how the flexible genomic region in RP62a populations evolved after 21 additional rounds of passaging in BHI (total of 50 passages; ∼333 generations). Sequencing of the P50 populations revealed that *S. epidermidis* underwent further genomic evolution. All five populations evolved additional deletions encompassing the anti-MGE defense island, with sizes ranging from ∼60 to ∼80 kb (Figure 6A). Two of the populations (A & B) exhibited a fixed deletion: population A lost all the anti-MGE systems and retained approximately 30% read coverage at the *mecA* gene, whereas population B lost both the *SCCmec* and anti-MGE systems (Figure 6A & B). In contrast, populations C, D, and E retained heterogeneous subpopulations carrying accessory genes, with detectable relative read depth for *mecA*,

**Figure 6.**
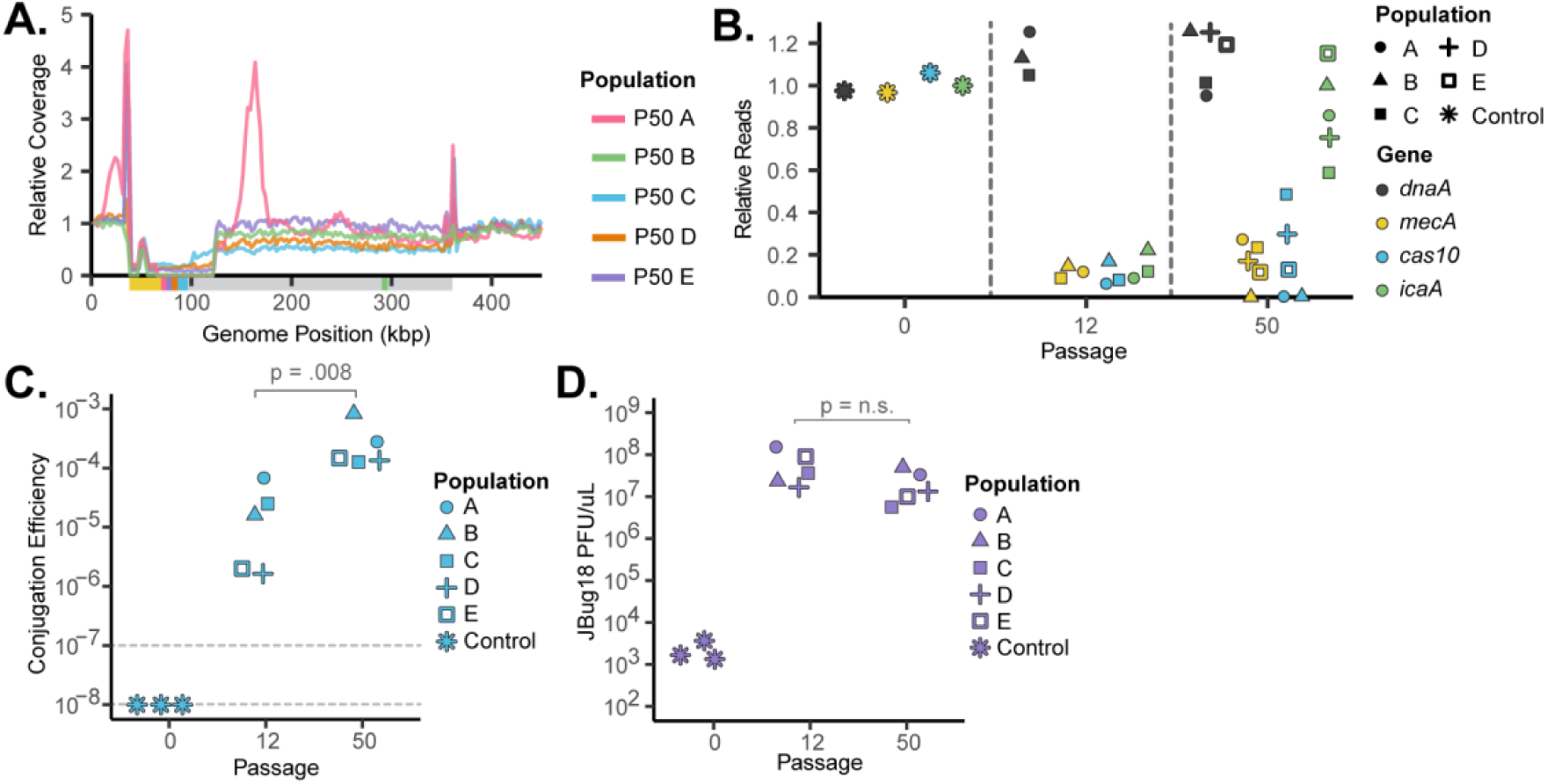
*S. epidermidis* populations show further genomic diversification at passage 50. **A.** Relative coverage plot of P50 populations replicates A-E. The x-axis shows the first 450 kbps out of 2.6 Mbps of the genome. The y-axis is the reads per million (RPM) normalized to the RPM at *dnaA* position. The colored region beneath the coverage plot is a schematic of the genes present in the region, as shown in Figure 1D. **B.** Relative reads of *dnaA, mecA, cas10,* and *icaA* from the sequencing data shown in Panel A **C.** pG0400 conjugation efficiency of the P0, P12, and P50 populations. The five evolved populations are indicated by the different shapes. **D.** Plaquing efficiency of the phage JBug18 at P0, P12, and P50. The five evolved populations are indicated by the different shapes.

*cas10*, and *icaA* (Figure 6A & B). Populations C and D also harbored the shorter deletions, while a subset of cells retained portions of the larger ∼320 kb region seen in P12 (Figure 6A& B). Notably however, the *icaA* genes appear to rebound in all populations by P50 (Figure 6B). Phenotypically, populations at P50 exhibited increased conjugation rates and phage sensitivity, mirroring the loss of the anti-MGE defenses at the genomic level (Figure 6C & D). Together, these data show that large genomic deletions targeting the anti-MGE defense island continue to evolve, with both the size of the deleted region and the degree of fixation varying across independently evolving populations. These results highlight the rapid rate with which *S. epidermidis* populations diversify.

## Discussion

Here, we show that evolving *S. epidermidis* populations can dynamically alter rates of HGT. Using experimental evolution, we found that *S. epidermidis* RP62a develops large genomic deletions of accessory genes, including an anti-MGE defense island, *SCCmec*, and the *ica* locus. Losing these genes increases HGT permissiveness, generates a heterogeneous β-lactam resistance phenotype, and reduces biofilm formation, respectively. In rich media, these deletions occur rapidly, within 80 generations, and are localized to a flexible genomic region downstream of *oriC*, resembling deletions observed in the *oriC* environ of related staphylococcal species, such as *S. haemolyticus* (Takeuchi et al., 2005). Mechanistically, the deletions are linked to IS elements, which can cause genomic deletions via transposases or homologous recombination (Albalat & Cañestro, 2016; Siguier et al., 2014; Wada et al., 1991). By approximately 333 generations, passaged populations show further diversification, with some developing shorter deletions that only remove the anti-MGE defense island. These later deletions could be facilitated by additional insertion elements in the flexible region or *ccrAB* attachment sites, which have been shown to mediate site specific excision events (Hossain et al., 2024).

A major consequence of the deletions is that *S. epidermidis* populations increase their ability to undergo HGT. In our results, evolved RP62a populations display orders-of-magnitude increases in their ability to accept plasmids via transformation and conjugation, at the cost of their susceptibility to lytic phage infection. This increase in HGT results from a loss of the anti-MGE defense island in the evolving *S. epidermidis* populations. Our work shows that these evolved heterogeneous populations can acquire and retain exogenous genes via HGT, such as new antibiotic resistance elements. These acquired genes, in turn, can improve the adaptability of the overall population to varying selective pressures compared to homogeneous populations. Because a subpopulation of the wild-type strain can be maintained, the population can acquire genetic novelty while retaining the beneficial genes on the accessory region, resulting in a population with composite phenotypes (Figure 7). Indeed, rapid evolution of heterogeneity has been reported in longitudinal studies on *S. epidermidis* isolates taken from human skin and is believed to be a survival strategy, allowing bacterial pathogens to adapt quickly to different environments (Datta et al., 2021; Schoenfelder et al., 2010; Severn & Horswill, 2023; Zhou et al., 2020).

**Figure 7.**
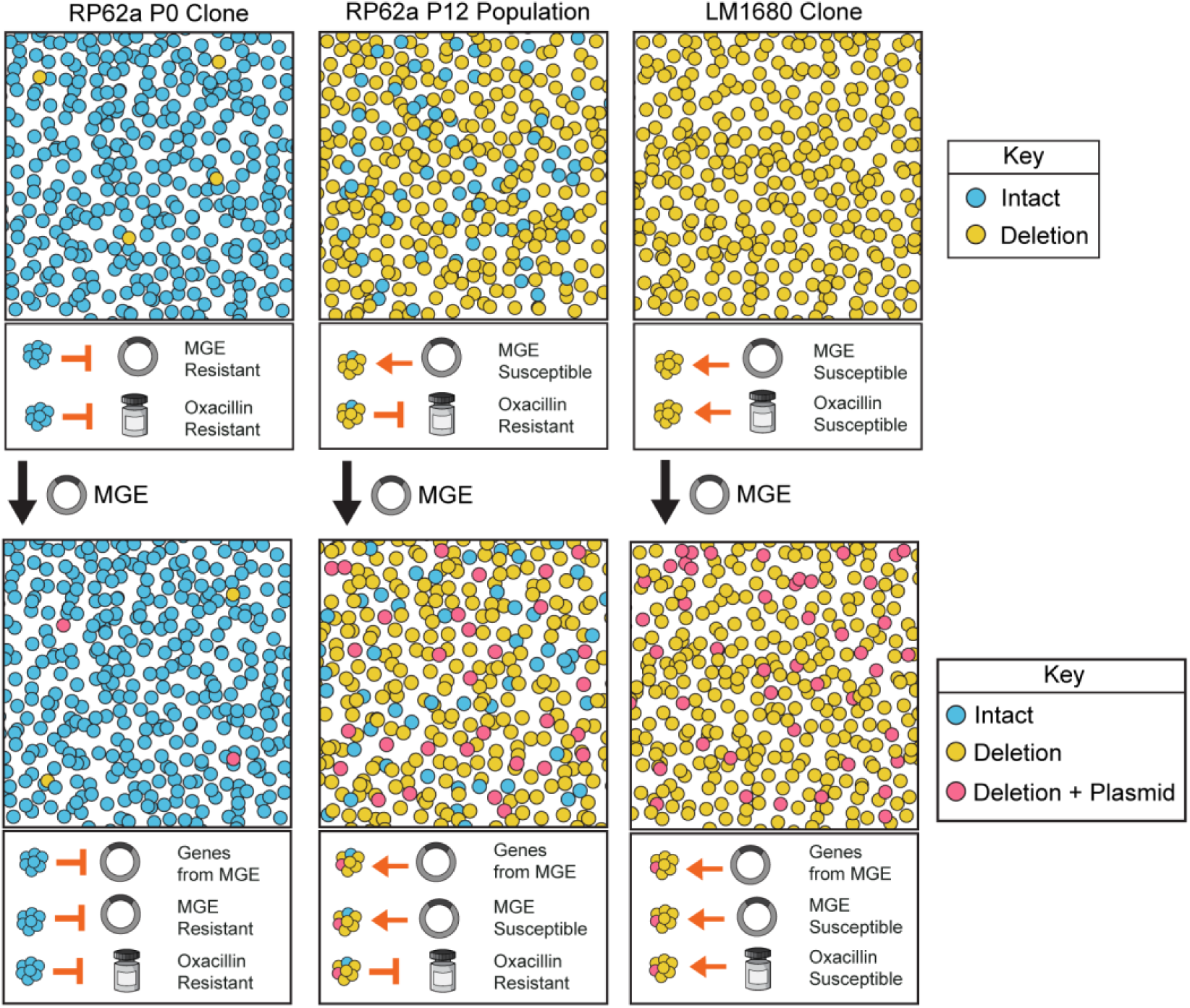
Heterogeneous anti-MGE defense in *S. epidermidis* populations improves HGT rates and antibiotic adaptability of the population. Homogenous wild-type RP62a populations are oxacillin resistant but prevent reduce the movement of MGEs, thereby limiting the acquisition of genes that can benefit survival (Left panel column). Homogenous LM1680 populations are oxacillin sensitive but can acquire genes through MGEs (Right panel column). Mixed P12 populations of RP62a remain oxacillin resistant but are permissive to HGT due to reduced anti-MGE defense. Acquisition of an antibiotic resistance gene can generate a population with composite resistance phenotypes.

Because anti-MGE genes and other accessory genes are clustered in a flexible genomic region, HGT levels can be modulated by applying different kinds of selective pressures to the population, such as antibiotic treatment or enrichment for sessile growth. These pressures stabilize the accessory gene region in the bacterial population, thereby retaining the anti-MGE islands and providing a means by which different selective pressures can indirectly influence the prevalence of defense systems within a population. This permits the maintenance of anti-MGE defense in the absence of selection exerted by MGEs (e.g. phages). Clustering of anti-MGE systems with other accessory genes is highly prevalent in *S. epidermidis*, *S. aureus,* and other bacterial species. We therefore speculate that this mechanism is applicable across staphylococci and that such selection-driven relationships are found in other bacterial species, such as *Vibrio cholerae,* where anti-MGE genes and resistance genes cluster (LeGault et al., 2021).

A limitation of this study is that these large-scale deletions were observed during growth in rich media conditions, which do not mimic the natural environments of *S. epidermidis*. We note, however, that patient studies have described heterogeneous *S. epidermidis* infections comprising of cells with ∼100 kB genomic deletions and cells with intact genomes. Notably, these populations became enriched for the intact 100-kb region during infection and antibiotic treatment (Weisser et al., 2010). These results suggest that our *in vitro* evolution experiments are potentially applicable to natural settings of *S. epidermidis.* Further, the linkage between anti-MGE elements, antibiotic resistance genes, and other accessory genes may need to be considered for combined phage/antibiotic therapies, since the application of one selective pressure can select for resistance to the other agent. Thus, *S. epidermidis* genomic plasticity and heterogeneity are key drivers in modulating HGT, allowing the population to protect against harmful MGEs while enabling the acquisition of beneficial MGEs. We conclude that environmental conditions and selective pressures outside of the anti-MGE defense systems can heavily influence HGT.

## Method

### Strains and Culture Conditions

The bacterial strains and phages used in this study are listed in Supplemental Table 1, along with the media conditions used. All bacterial cultures were grown at 37 °C with shaking (235 RPM), unless otherwise indicated. Phages were stored at 4 °C.

### Passaging conditions

#### Rich media conditions

Bacterial clones were picked and inoculated into 5 mL of BHI (Brain Heart Infusion) media supplemented with erythromycin (Erm; 10 µg/mL) and neomycin (Neo; 15 µg/mL). Cultures were grown at 37 °C with shaking. Starting at P0, each replicate culture was diluted 1:100 into tubes containing 5 mL of fresh BHI Erm10/Neo15 (population A- E). The cultures were passaged 1:100 every 24 hours into fresh BHI Erm10/Neo15 media. 50% glycerol stocks were prepared every 4 passages up to Passage 50 and stored at −80 °C.

#### Passaging under oxacillin selection

Five P0 clones (Replicates A-E) were picked and inoculated into 5 mL of BHI Erm10/ Neo15. Cultures were grown at 37 °C with shaking. Each of the replicates was passaged every 24 hours 1:100 into fresh 5 mL of BHI Erm10/Neo15 supplemented with oxacillin at concentrations of 12 µg/mL, 6 µg/mL, 3 µg/mL, and 0 µg/mL. Glycerol stocks were prepared every 6 passages, up to Passage 15.

#### Passaging under sessile conditions

Protocol modified from Wiebuhr et. al. 1999. 20 mL of Tryptic Soy Broth (TSB) supplemented with erythromycin (Erm; 10 µg/mL) and neomycin (Neo; 15 µg/mL) was added to a polystyrene tissue culture flask and was inoculated with a bacterial culture from glycerol. The culture was incubated under static conditions at 37 °C. After 24 hours, the medium was replaced. The procedure was repeated for 15 passages. The culture was washed with PBS, and adherent cells were scraped and made into glycerol stocks for further phenotypic analysis.

### Treatment of evolved populations with various selective pressures

#### Treatment with oxacillin

RP62a P12 populations were grown directly from glycerol stocks. The bacterial culture was diluted 1:100 and incubated at 37 °C with shaking until the population was in mid-log phase (OD_600_ = 0.3). 12 μg/mL of oxacillin was added, and the population was incubated at 37 °C with shaking for 18 hours. The treated culture was then preserved as a glycerol stock for further analysis.

#### Treatment with SP

The evolved populations were inoculated in a polystyrene tissue culture flask containing 20 mL of TSB Erm10/Neo15. The protocol follows the passaging under sessile conditions, except that the procedure was repeated for 9 passages.

#### Treatment with phage infection

Overnight cultures of RP62a P12 populations were grown directly from glycerol stock. The overnight culture was diluted 1:100 into 5 mL of BHI Erm10/Neo15, allowed to enter mid-log phase (OD_600_ = 0.3) at 37 °C with shaking, and infected with phage at an MOI of 1 (unless noted otherwise). The treated culture was grown at 37 °C under shaking conditions. The treated culture was then preserved as a glycerol stock.

### Alternating treatment of evolved populations with antibiotics (R1→R2)

RP62a P12, RP62a P0, and LM1680 were grown directly from glycerol stocks. For round 1 (R1), the bacterial cultures were diluted 1:100 into BHI and incubated at 37 °C with shaking until the population was in mid-log phase (OD_600_ = 0.3), after which either 12 µg/mL of oxacillin or 5 µg/mL mupirocin was added. Populations were incubated at 37 °C with shaking for 18 hours. A portion of treated culture was tested for its ability to grow under increasing amounts of oxacillin or mupirocin in a plate reader-based growth assay (see ‘Antibiotic growth assay**’** below). For round 2 (R2), a portion of the bacterial cultures was diluted 1:100 into BHI and incubated at 37 °C with shaking until the population was in mid-log phase (OD_600_ = 0.3). The second antibiotic was then added. Populations were incubated at 37 °C with shaking for 18 hours, after which populations were then tested for their growth under increasing amounts of oxacillin or mupirocin.

### DNA Sequencing and Genome Assembly

Phenol-chloroform extraction was performed to isolate genomic DNA from passaged bacterial populations and individual clones. The purified gDNA from the population and individual clones was sent to SeqCenter for Illumina sequencing, and to Plasmidsaurus for Oxford Nanopore sequencing. Genomic assembly was performed in Geneious Prime (version 2026.0.2) using S. epidermidis RP62a (RefSeq: NC_002976.3) as the reference genome (Supplemental Table 1).

### Data Visualization and Statistical Analysis

Data visualizations and statistical analyses using R v4.3.2, primarily with the tidyverse package.

### Relative Reads and Genomic Coverage

Post-genomic assembly, Geneious was used to generate a coverage table that showed the number of reads at each genomic position. To normalize the coverage, we calculated the reads per million (RPM). RPM is calculated using the following formula: number of sequences at a genomic position divided by the total number of sequences, then multiplied by 10^6^. We binned the RPM for every 2.5 kbp. This normalization method had varying reads of *dnaA* across different genomic sequences. To facilitate comparison across different genomic sequences, we calculated relative coverage by normalizing RPM using the following formula: RPM divided by RPM for *dnaA*. The relative reads were determined by dividing the number of reads at a specific position by the average total number of reads. We calculated the relative reads for the genes *dnaA*, *mecA*, *cas10*, and *ica*.

### Phage Propagation

Overnight cultures of LM1680 were diluted 1:100, grown to mid-log phase (OD_600_ = 0.3), treated with phage, and incubated for 4 hours. Centrifugation and filter sterilization were performed with a 0.22 μm syringe filter to purify phage particles. Phage titer was quantified using a double overlay plaque assay.

### Antibiotic growth assay

For the antibiotic MIC agar overlay assays, MIC strips were placed on bacterial lawns prepared by mixing 1×10^8^ CFU/mL with 5 mL of 0.5x BHI top agar and overlaying the mixture onto BHI agar plates. The plates were incubated at 37°C for 18 hours. For growth in liquid culture, BHI with the antibiotic was serially diluted in 2-fold increments on a 96-well plate. The plates were inoculated with 1×10^5^ CFU/mL of the bacterial culture. Using a Tecan plate reader, OD_600_ was measured every 10 minutes for 24 hours or as noted. The plate was shaking at 37 C.

### Congo Red Assay

BHI Congo red agar was modified from Ziebuhr et. al., 1999 and Freeman et. al., 1989 5/1/2026 3:16:00 PM, with 5 g/L sucrose and 0.8 g/L Congo red dye added to a BHI agar mixture. 4 µL of the overnight culture was spotted onto the Congo red plate. The plates were incubated at 37°C for 18 hours. The plates were left at room temperature for another 24 hours before quantification. Black colonies suggest PIA^+^ and red colonies suggest PIA(-) (Freeman et al., 1989; Ziebuhr et al., 1999).

### Plate-Based Plaque Assay

Bacterial lawns were prepared by mixing 100 µL of overnight culture with 5 mL of 0.5x melted BHI agar. The mixture was poured onto a solid BHI erm10 neo15 plate and allowed to dry for 10 minutes. Phages were serially diluted 10-fold (10^-1^-10^-8^ unless otherwise noted), and 3 μL of each dilution was spotted onto a bacterial lawn. The plates were then incubated at 37 °C for 18 hours. Plaque-forming units per μL were quantified.

### Conjugation Assay

Overnight cultures of the donor and recipient were diluted 1:100 in 5 mL of BHI and grown at 37 °C with shaking at 235 RPM until the OD reached 0.8. The donor and recipient were then mixed at a 1:1 ratio (500 μL of each culture). The mixed culture was then pelleted, washed with BHI, and subjected to another pellet step. The culture was then resuspended in 100 μL of BHI and plated onto a BHI agar plate using a 0.45 μm filter paper. The plate was then incubated at 37 °C for 18 hours. Using forceps, the filter paper containing the mixed bacterial culture was resuspended in 5 mL of liquid BHI. The culture was then serially diluted 10-fold (10^0^-10^-7^) in liquid BHI. 5 μL of the serially diluted culture was spotted and allowed to drip onto both selective and non-selective plates. Selective plates consisted of BHI agar with Erm10, Neo15, and Mupirocin (Mup; 5 µg/mL); non-selective plates consisted of BHI agar with Erm10/Neo15. Plates were incubated at 37 °C without shaking for 18 hours. The plates were enumerated by calculating the CFU. Conjugation efficiency was calculated as a ratio of CFU between conjugants and transconjugants. The limit of detection (LOD) is the lowest ratio that can be detected but cannot be accurately estimated. The limit of detection is the ratio of the minimum cell count (1 CFU) divided by the maximum cell count (10^-8^ CFU).

### Transformation Assay

*S. epidermidis* populations were made electrocompetent with established methods and electroporated with normalized levels of pC194 with a chloramphenicol resistance marker (J. Y. H. Lee et al., 2019b). The cells were resuspended in BHI and allowed to recover in the 37 °C incubator at 235 RPM for 2 hours. The cultures were serially diluted 10-fold, and 100 µL of each dilution was plated on selective agar plates (BHI agar Erm10/Neo15/Cm10) and non-selective plates (BHI Erm10/Neo15). Transformation efficiency was calculated by dividing the colony-forming units on selective by non-selective plates.

### Competition experiments between LM1680 and RP62a

Plasmid pC194 was transformed into LM1680 to confer chloramphenicol resistance. Overnight clonal cultures of LM1680 pC194 and RP62a were incubated overnight at 37 °C at 235 RPM. After incubation, optical density (OD) was measured at 600 nm for both cultures, and the bacterial density was normalized to the lower OD. Varying ratios of RP62a and LM1680 pC194 (1:1, 1:1000, 1000:1) were passaged at 1:100 into fresh 5 mL BHI Erm10/Neo15. The culture was incubated overnight at 37 °C at 235 RPM. Following incubation, the cultures were passaged 1:100 into fresh 5 mL BHI Erm10/Neo15, and the process was repeated for 7 passages. To quantify CFUs at each passage, the population was plated onto BHI agar with Neo15/Erm10/Cm10 (to select for LM1680) and Neo15/Erm10/Oxi2 (to select for RP62a). Relative abundance of LM1680 pC194 (A) and RP62a (B) at varying initial ratios through 7 passages was calculated as A/(A+B) and B/(A+B). The change in relative fitness was calculated with the formula Δr = ln(RP62a_final_/RP62a_initial_) − ln(LM1680_final_/LM1680_inital_) per passage intervals.

### Identification of defense systems and insertion sites (IS) across *S. epidermidis* genomes

To assess the prevalence of IS elements across *Staphylococcus epidermidis* genomes, we retrieved all complete *S. epidermidis* genomes from NCBI using the ncbi-dataset command line tools (datasets download genome taxon ‘Staphylococcus epidermidis’ --assembly-level complete --include genome,cds,protein,gbff --mag exclude --assembly-source ‘RefSeq’). This resulted in 225 genomes, including the reference genome of RP62a. Next, a Python script to extract the sections of each genome from the gene (*dnaA*) to 1,000,000 bp (1 million) bp downstream of the *rlmH* gene based on the information in the gene annotation file (GFF file), resulting in 201 genomes. The 201 genomes segments (*dnaA* to 1Mbp downstream) were concatenated into 1 fasta file, and used to create the BLAST database. IS431mec mediates the excision of the accessory region. To identify IS431 across bacterial genomes, we used the reference IS431-meclike from ISFinder and performed a blastn search. Only results with an identity greater than 98% and e-values ranging from 0 to 4.01e-22 were retained for plotting in R using the tidyverse package. Next, we classified a broader range of IS elements using the software ISEScan (v.1.7.3). Only results for which the IS element was classified as “complete” were kept for plotting (Figure 1B). Finally, defense and anti-defense systems were found in the 201 genomes by using the software defenseFinder (v.2.0.1). Code availability : https://github.com/MolabUW/staphepi_IS

### Identification of mutations across *S. epidermidis* genomes

Sequencing reads were filtered with fastp (version 1.3.1) with a quality score cutoff of 28 and the --dedup and --trim_poly_g options. The genomic samples were then aligned to the latest *S. epidermidis* RP62a reference genome. Mutations in each sample were identified using breseq (version 0.38.1)(Deatherage & Barrick, 2014). Mutations identified in the ancestral P0 clone were excluded prior to subsequent analyses.

## Acknowledgements

MT is supported by the SciMed GRS program at UW-Madison; CYM is supported by start-up funds from the Department of Bacteriology at UW-Madison and the Margaret Q. Landenberger Foundation. We thank the Marraffini and Hatoum-Aslan laboratories for providing us with bacteriophage samples

## Supplemental Figures

**Supplemental Figure 1:**
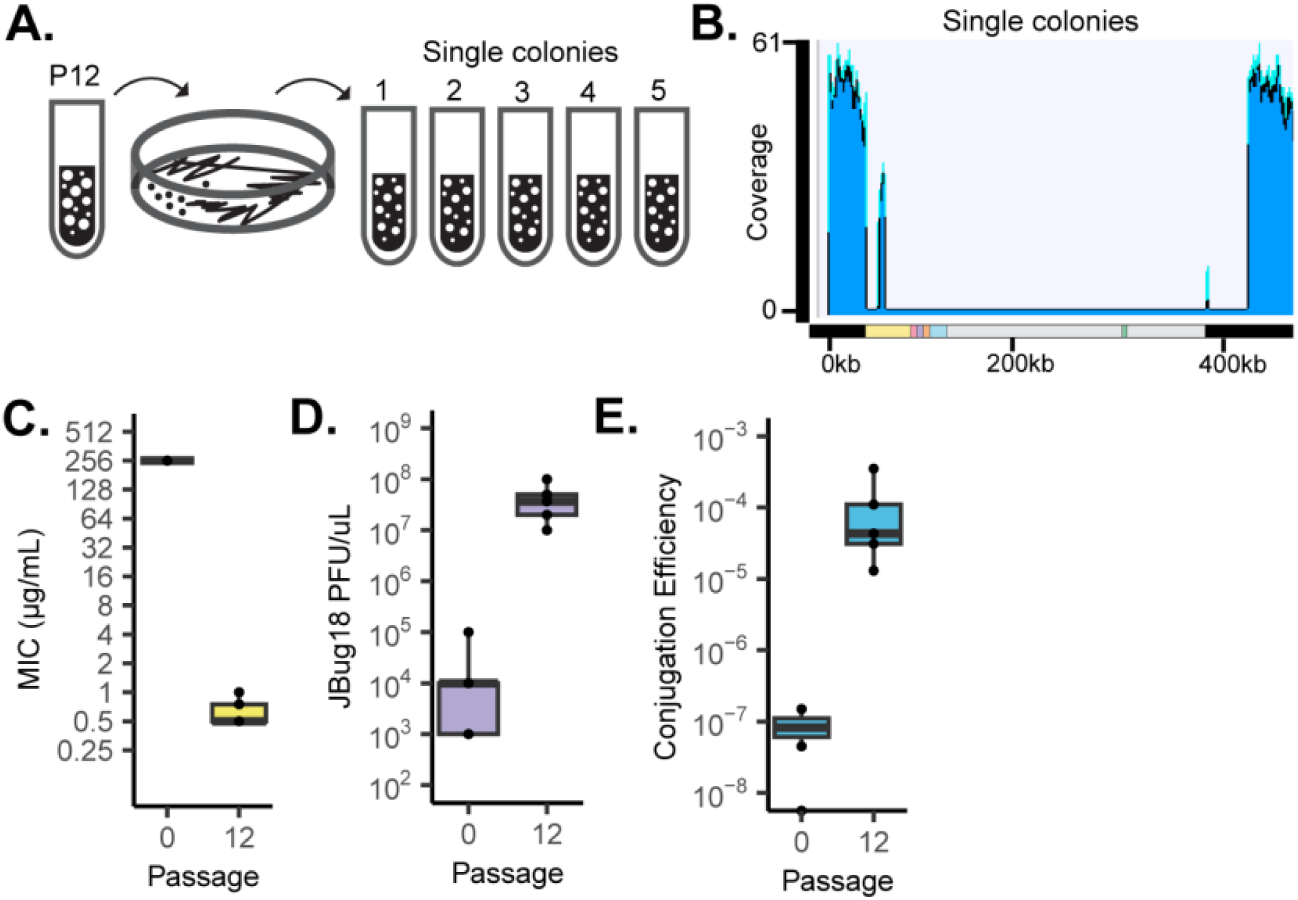
Clones from the RP62a P12 population exhibit a large-scale deletion. **A.** Cells of RP62a P12 were streaked onto BHI plates, and 5 clones were picked from the plate for further testing. **B**. Genomic coverage of RP62a P12 clone A exhibits a deletion size of ∼320 kB. Sequencing results were obtained from Oxford Nanopore sequencing. **C-E.** Phenotypic assays on the 5 evolved RP62a clones compared to the parental clone P0. Box plots represent data from each independent clone. The boxes indicate the interquartile range, the horizontal lines indicate the median, and the whiskers represent the minimum and maximum values. **C.** Minimum Inhibitory Concentration (MIC) of the clones against oxacillin. **D.** Plaquing efficiency of the bacteriophage JBug18. **E.** Conjugation efficiency of the staphylococcal plasmid pG0400.

**Supplemental Figure 2:**
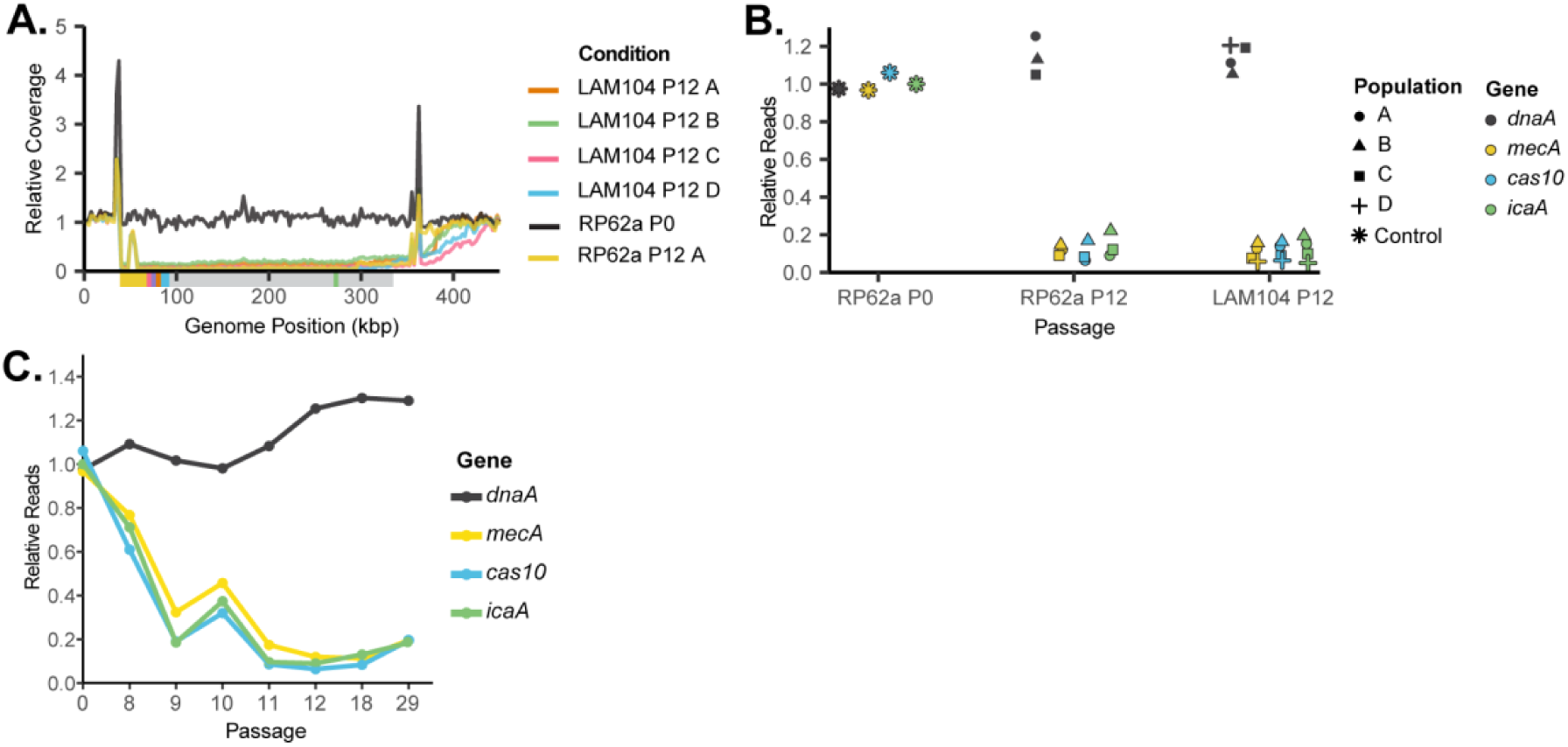
Dynamics of the genomic deletion in *S. epidermidis* RP62a and LAM104. **A.** Relative read coverages of the evolved LAM104 populations grown in rich media (BHI). The x-axis is the first 450 kb out of 2.6 Mbp of the genome. The y-axis is the reads per million (RPM) normalized to the RPM at *dnaA* position. **B.** Relative reads of the evolved LAM104 population compared to the wild-type strains RP62a. Shown are the relative abundances of *dnaA, mecA, cas10,* and *icaA*. **C.** Relative reads measuring the presence of a gene in the population through passages. Relative reads were measured by dividing the average number of reads at a gene locus by the average number of total reads. The relative reads were calculated across P0 and P8-P29 for Population A.

**Supplemental Figure 3:**
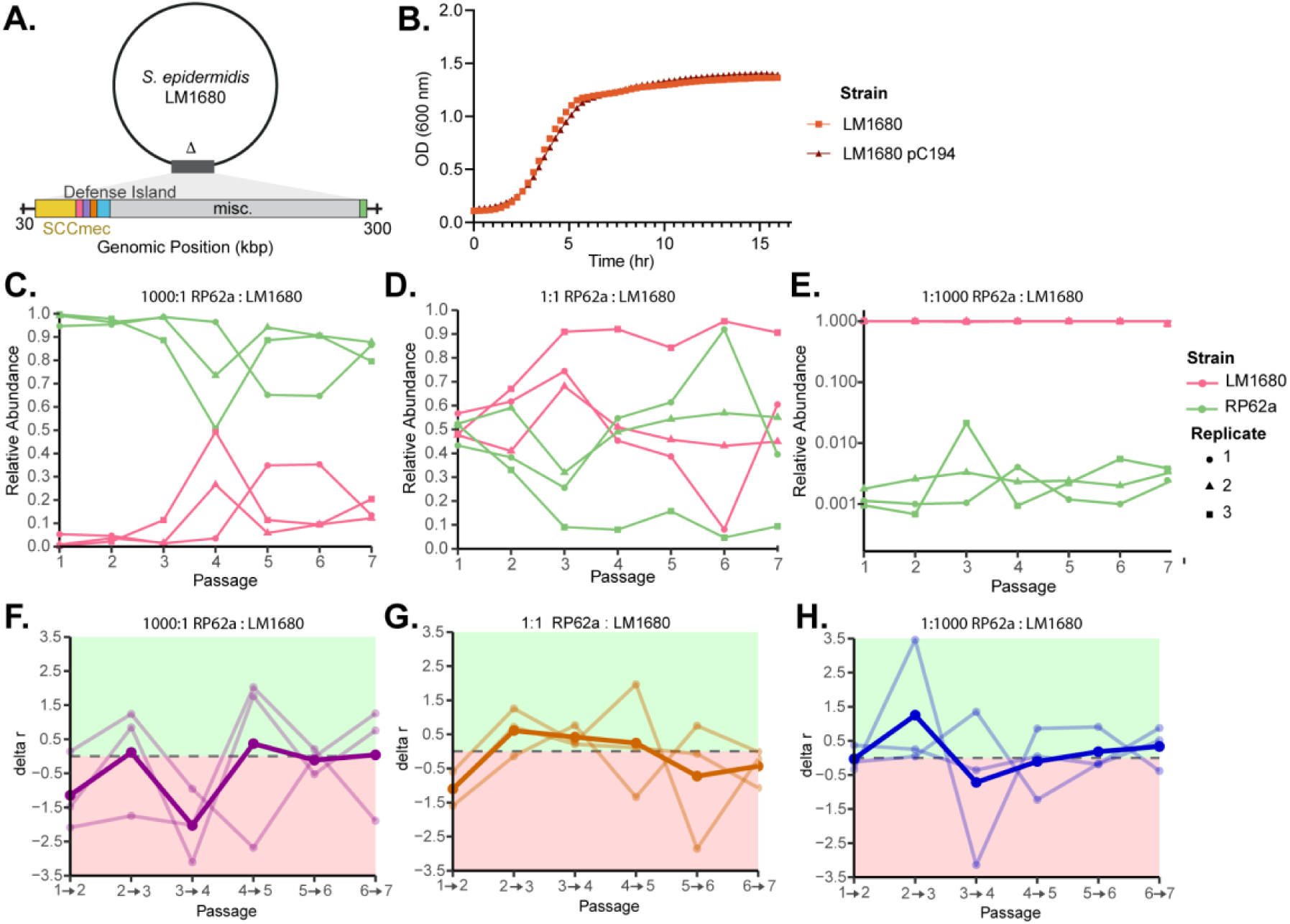
Competition experiments between wild-type *S. epidermidis* RP62a and LM1680 (Daccessory) **A.** Genome of *S. epidermidis* LM1680 lacks *SCCmec*, the anti-MGE defense island, and *ica* genes. **B.** Growth curves of LM1680 with and without the staphylococcal plasmid pC194. Error bars represent the standard deviation of 3 biological replicates. **C-E.** Relative abundance of passaged co-cultures of LM1680 + pC194 (red) RP62a (green). RP62a and LM1680 were mixed at starting ratios of (**C)** 1000:1, (**D)** 1:1, and (**E)** 1:1000. Each line represents an independent biological replicate. **F-H.** Relative fitness of passaged co-cultures of LM1680 + pC194 and RP62a. Change in fitness rate (Δr = ln(RP62a_final_/RP62a_initial_) − ln(LM1680_final_/LM1680_inital_) of RP62a and LM1680 per passage intervals. Δr > 0 indicates a growth advantage for RP62a (green); Δr < 0 indicates a growth advantage of LM1680 (red); Δr = 0 indicates neutral growth, indicated by a dashed line. Bold line indicates the mean of n = 3 independent populations. Lighter lines indicate the independent populations. Relative fitness of LM1680:RP62a populations mixed at (**F)** 1000:1, (**G)** 1:1, and (**H)** 1:1000 ratio.

**Supplemental Figure 4:**
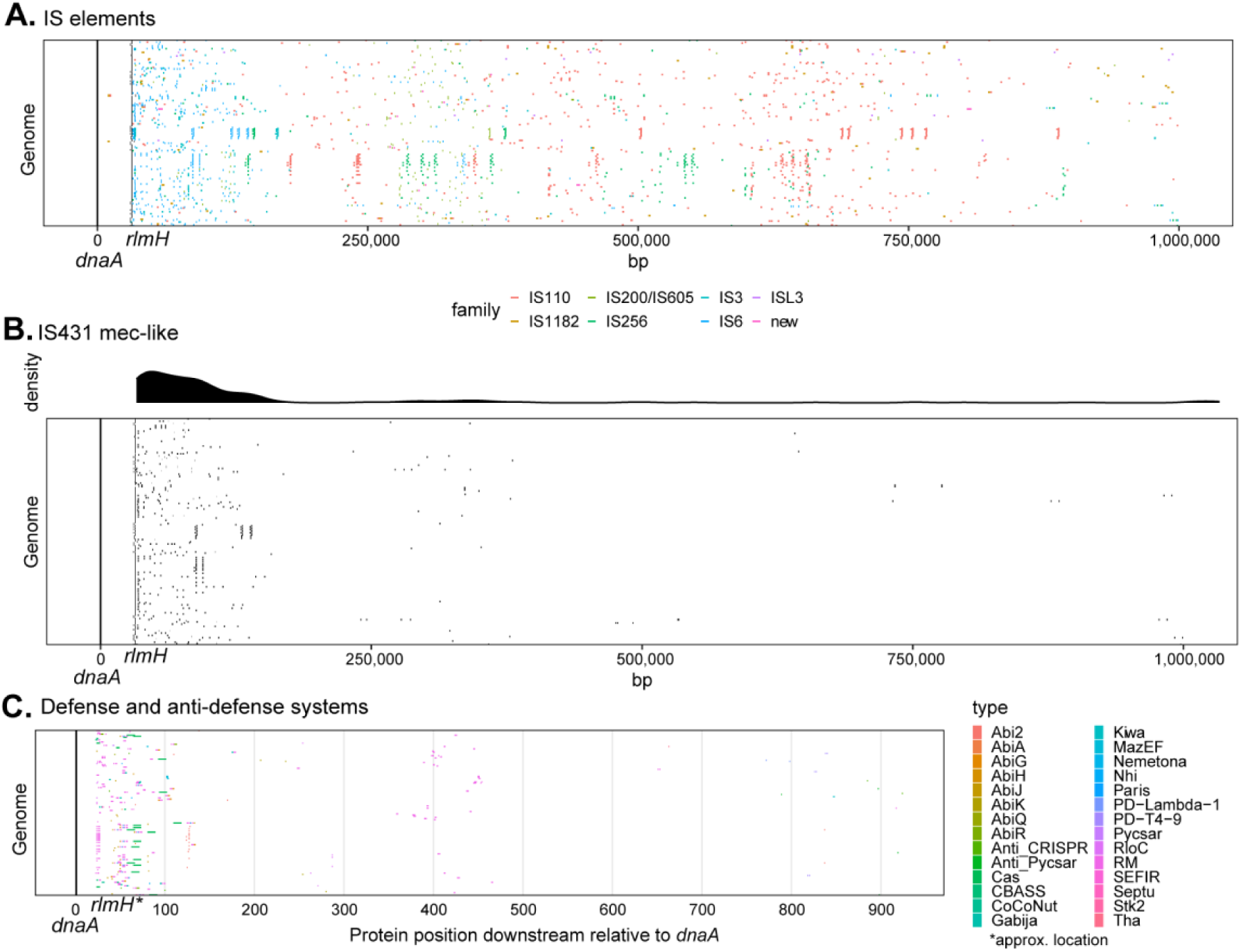
IS elements cluster near anti-MGE systems. **A.** Putative IS elements found across 201 *S. epidermidis* genomes. **B.** IS431-mec distribution across 201 of *S. epidermidis* genomes. **C.** Defense and anti-defense systems predicted across 201 *S. epidermidis* genomes. IS elements (Panel A,B) were found using ISFinder and ISEScan; defense systems (Panel C) were found using DefenseFinder.

**Supplemental Figure 5:**
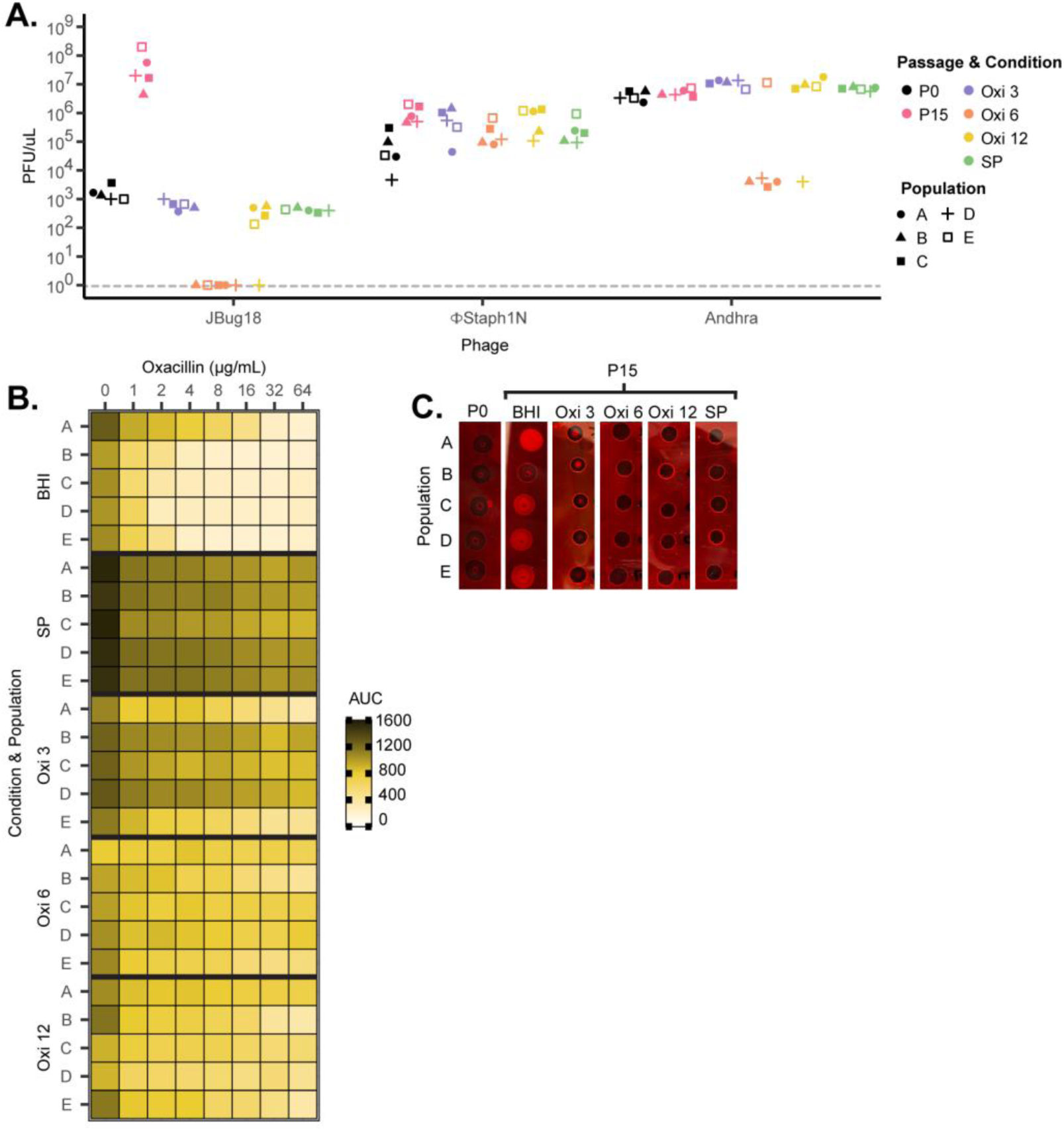
Passaging RP62a with oxacillin or in a static environment selects for the maintenance of the accessory genes at a population level. **A.** Plaque titer (PFU/µL) of phages JBug18, FStaph1N, and Andhra on bacterial background of *S. epidermidis* populations P0, P15 (non-selective media), Oxi 3, Oxi 6, and Oxi 12 (Passage 15 in the presence of 3 µg/mL, 6 µg/mL, and 12 µg/mL of oxacillin, respectively), and SP. Individual replicates A-E represented by distinct shapes. **B.** Heatmap showing the area under the curve for P15 *S. epidermidis* populations grown under increasing levels (µg/mL) of oxacillin. Individual replicates are indicated as A-E. **C.** Congo Red assays on *S. epidermidis* populations P0, P15, Oxi 3, Oxi 6, Oxi 12, and SP. Black coloration indicates PIA(+), while red coloration indicates PIA(-).

**Supplemental Figure 6:**
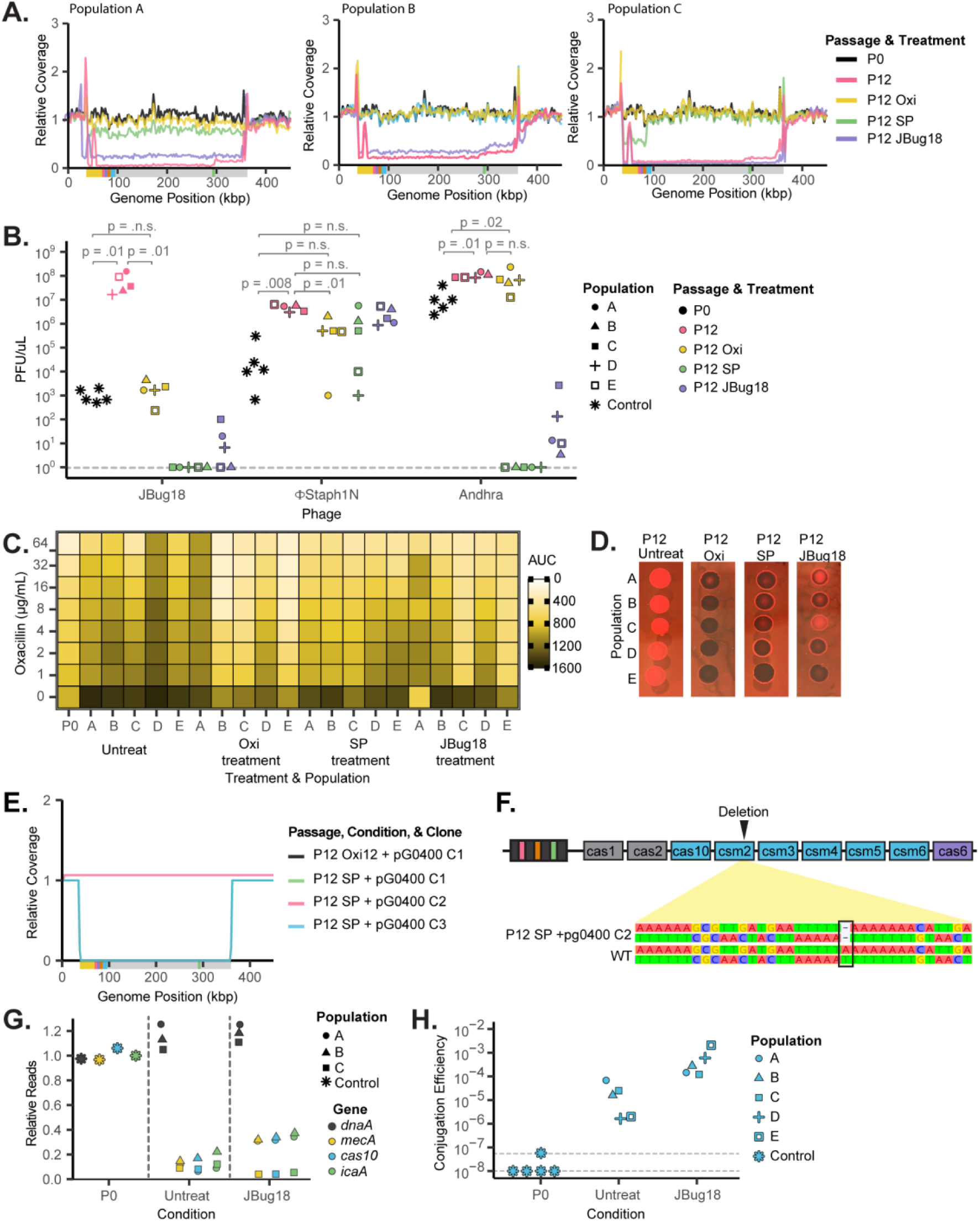
Genotypic and phenotypic analyses of evolved S. epidermidis populations treated with oxacillin, sessile passaging, or phage infection. **A-C.** Relative read coverages of three independent evolved *S. epidermidis* RP62a populations treated with oxacillin (P12 Oxi), infected JBug18 phage (P12 JBug18), or passaged in sessile populations (P12 SP). The x-axis is the first 450 kb out of 2.6 Mbp of the genome. The y-axis is the reads per million (RPM) normalized to the RPM at *dnaA* position. Coverage plots of untreated P0 and P12 populations are shown as controls. **B.** Phage titers of (PFU/µL) of JBug18, Andhra, and FStaph1N on treated and untreated P12 populations. Population conditions are the same as in Panel A. Individual replicates A-E are represented by distinct shapes. **C.** Heatmap showing the area under the growth curve (AUC) for treated and untreated *S. epidermidis* P12 populations grown under increasing levels (µg/mL) of oxacillin. Population conditions are the same as in Panel A. Individual replicates are indicated as A-E. **D.** Congo Red assays on treated and untreated *S. epidermidis* populations. Population conditions are the same as in Panel A. Black coloration indicates PIA(+), while red coloration indicates PIA(-). **E.** Long-read sequencing data of treated (Oxi12 or SP) *S. epidermidis* clones that acquired the pG0400 plasmid. **F.** Frameshift mutation in the *csm2* gene of the CRISPR locus of P12 SP + pG0400 C2. **G.** Relative abundances of *dnaA, mecA, cas10,* and *icaA* in JBug18-treated *S. epidermidis* P12 populations. **H.** pG0400 conjugation efficiency of JBug18-treated *S. epidermidis* P12 populations.

